# BA.2.12.1, BA.4 and BA.5 escape antibodies elicited by Omicron infection

**DOI:** 10.1101/2022.04.30.489997

**Authors:** Yunlong Cao, Ayijiang Yisimayi, Fanchong Jian, Weiliang Song, Tianhe Xiao, Lei Wang, Shuo Du, Jing Wang, Qianqian Li, Xiaosu Chen, Yuanling Yu, Peng Wang, Zhiying Zhang, Pulan Liu, Ran An, Xiaohua Hao, Yao Wang, Jing Wang, Rui Feng, Haiyan Sun, Lijuan Zhao, Wen Zhang, Dong Zhao, Jiang Zheng, Lingling Yu, Can Li, Na Zhang, Rui Wang, Xiao Niu, Sijie Yang, Xuetao Song, Yangyang Chai, Ye Hu, Yansong Shi, Linlin Zheng, Zhiqiang Li, Qingqing Gu, Fei Shao, Weijin Huang, Ronghua Jin, Zhongyang Shen, Youchun Wang, Xiangxi Wang, Junyu Xiao, Xiaoliang Sunney Xie

## Abstract

SARS-CoV-2 Omicron sublineages BA.2.12.1, BA.4 and BA.5 exhibit higher transmissibility over BA.2^1^. The new variants’ receptor binding and immune evasion capability require immediate investigation. Here, coupled with Spike structural comparisons, we show that BA.2.12.1 and BA.4/BA.5 exhibit comparable ACE2-binding affinities to BA.2. Importantly, BA.2.12.1 and BA.4/BA.5 display stronger neutralization evasion than BA.2 against the plasma from 3-dose vaccination and, most strikingly, from post-vaccination BA.1 infections. To delineate the underlying antibody evasion mechanism, we determined the escaping mutation profiles^2^, epitope distribution^3^ and Omicron neutralization efficacy of 1640 RBD-directed neutralizing antibodies (NAbs), including 614 isolated from BA.1 convalescents. Interestingly, post-vaccination BA.1 infection mainly recalls wildtype-induced humoral memory. The resulting elicited antibodies could neutralize both wildtype and BA.1 and are enriched on non-ACE2-competing epitopes. However, most of these cross-reactive NAbs are heavily escaped by L452Q, L452R and F486V. BA.1 infection can also induce new clones of BA.1-specific antibodies that potently neutralize BA.1; nevertheless, these NAbs are largely escaped by BA.2/BA.4/BA.5 due to D405N and F486V, and react weakly to pre-Omicron variants, exhibiting poor neutralization breadths. As for therapeutic NAbs, Bebtelovimab^4^ and Cilgavimab^5^ can effectively neutralize BA.2.12.1 and BA.4/BA.5, while the S371F, D405N and R408S mutations would undermine most broad sarbecovirus NAbs. Together, our results indicate that Omicron may evolve mutations to evade the humoral immunity elicited by BA.1 infection, suggesting that BA.1-derived vaccine boosters may not achieve broad-spectrum protection against new Omicron variants.

## Main

The recent emergence and global spreading of severe acute respiratory syndrome coronavirus 2 (SARS-CoV-2) variant Omicron (B.1.1.529) have posed a critical challenge to the efficacy of COVID-19 vaccines and neutralizing antibody therapeutic^6–9^. Due to multiple mutations to the spike protein, including its receptor-binding domain (RBD) and N-terminal domain (NTD), Omicron BA.1 can cause severe neutralizing antibody evasion^3, 10–13^. Currently, Omicron sublineage BA.2 has rapidly surged worldwide, out-competing BA.1. Compared to the RBD of BA.1, BA.2 contains three additional mutations, including T376A, D405N and R408S, and lacks the G446S and G496S harbored by BA.1 (Extended Data Fig. 1a). The S371L on BA.1 is also substituted with S371F in BA.2. Importantly, new Omicron variants are still continuously emerging. The recently appeared new Omicron variants contain identical RBD sequences to BA.2 but with the addition of L452 and F486 substitutions, namely BA.2.12.1 (L452Q), BA.2.13 (L452M), BA.4 and BA.5 (L452R+F486V), and all displayed higher transmission advantage over BA.2. The new variants’ receptor binding and immune evasion capability require immediate investigation.

### Structural analyses of Omicron Spike

First, we expressed and purified the prefusion-stabilized trimeric ectodomains of BA.1, BA.2, BA.3, BA.2.12.1, BA.2.13 and BA.4/BA.5 Spike (S-trimer). Noteworthy, BA.4 and BA.5 share the same spike mutations. All spike-trimers contain GSAS and 6P mutations along with the T4 fibritin trimerization domain for stabilization purpose^14, 15^. We determined the cryo-EM reconstructions of these S-trimers at an overall resolution of 3.1-3.5 Å, together with our previous reported BA.1 structure, allowing us to compare the detailed structural variations across Omicron sublineages (Fig. 1a and Extended Data Fig. 1b). Distinct from stably maintaining an open conformation with one ‘up’ RBD and two ‘down’ RBDs observed in BA.1 S-trimer ^16^, BA.2 and BA.2.12.1 exhibit two conformational states corresponding to a closed-form with all three RBDs “down” and an open form with one RBD “up”. Notably, one RBD was clearly disordered, representing a stochastic movement in BA.2.13, which, together with BA.2 and BA.2.12.1, suggests structural heterogeneity in the S-trimers of BA.2 sublineages. Surprisingly, most BA.3 and BA.4 S-trimers adopt closed- or semi-closed forms (Fig. 1a). The RBD confirmation differences could be allosterically modulated by mutations/deletions in NTD or near the furin cleavage site, with the detailed mechanism unclear. Also, the BA.4/5 spike we used contains the N658S mutation, which was presented in early BA.4/5 sequences but later disappeared due to a disadvantage in transmissibility, and may correlate with BA.4/5’s more closed RBD configurations. Interestingly, BA.2 sublineage S-trimers harbor relatively less compacted architectures in the region formed by the three copies of S2 (Fig. 1b). By contrast, BA.1, BA.3 and BA.4/BA.5 possess relatively tight inter-subunit organization with the more buried areas between S2 subunits (Fig. 1b). In line with structural observations, thermal stability assays also verified that S-trimers from BA.2 sublineages were the least stable among these variants, which might confer enhanced fusion efficiencies (Fig. 1c).

**Fig. 1.**
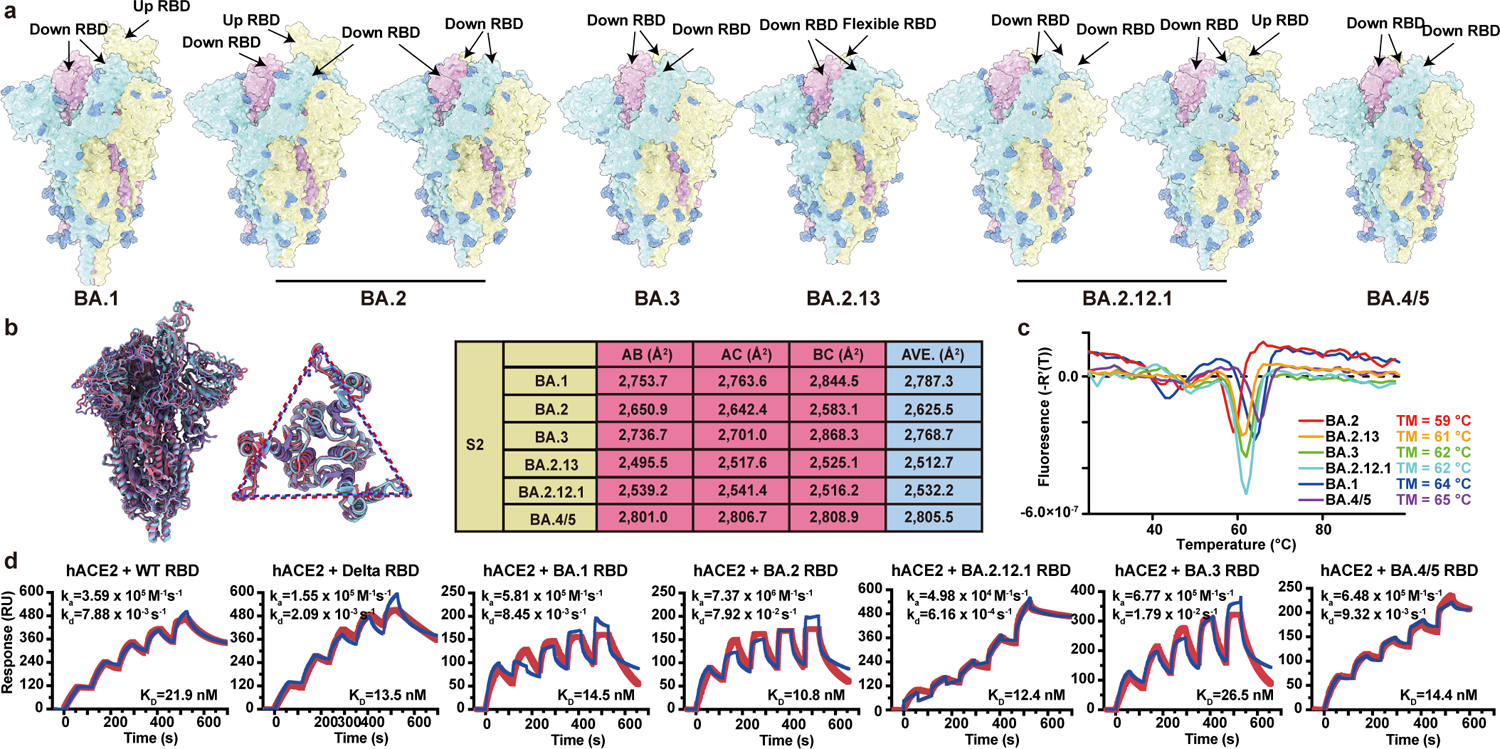
Structural and receptor-binding characteristics of Omicron subvariants. **a**, Surface representation for structures of S trimer of BA.1, BA.2, BA.3, BA.2.13, BA.2.12.1 and BA.4/5 variants. **b**, Structural interpretation and functional verification for the stability of S protein for BA.1, BA.2, BA.3, BA.2.13, BA.2.12.1 and BA.4/5 variants. The superimposed structures for S protein and the S2 domains of BA.1 (purple), BA.2 (red) and BA.4/5 (blue) are shown on the left. The binding surface areas between S2 subunits of all variants are calculated in the table on the right. **c**, Thermoflour analysis for these Omicron variants. Curves for each Omicron variant are colored in rainbow (BA.1: blue; BA.2: red; BA.3: green; BA.2.13: orange; BA.2.12.1: cyan; BA.4/5: purple). Thermoflour analyses were conducted in biological duplicates. **d**, Binding affinities of RBDs of Omicron variants to hACE2 measured by SPR. SPR analyses were conducted in biological duplicates.

Next, we measured the binding affinity between hACE2 and S-trimers of the Omicron variants by surface plasmon resonance (SPR) (Extended Data Fig. 1c). BA.4/5 spike trimer showed a decreased hACE2 binding affinity than other Omicron subvariants; however, this measurement could be deficient due to the additional N658S mutation. To exclude N658S’s potential influence, the binding affinities between hACE2 and Omicron variants’ RBDs are also examined (Fig. 1d). The RBD of Delta and all circulating Omicron subvariants exhibited similar ACE2 binding affinity, except for BA.3, which showed lower affinity, comparable to that of the wildtype (WT) strain. Additionally, the BA.2 subvariants displayed slightly increased hACE2 binding affinities than other Omicron variants. To further explore the molecular basis for altered binding affinities of these variants to hACE2, we performed molecular dynamics (MD) simulations based on cryo-EM structures and examined the effect of the RBD residue substitutions on the interaction with hACE2 (Extended Data Fig. 1d). Results revealed that the lack of G496S in BA.2 sublineages retained the hydrogen bond with K353 to hACE2, increasing their binding capability, which is in line with experimental observations revealed by deep mutational scanning assay^17^. Unexpectedly, a local conformational perturbation surrounding residues 444-448 lost its hydrophilic interaction between S446 with Q42 from hACE2 in BA.3, which is presumably caused by the single mutation G446S rather than double mutations of G446S and G496S (Extended Data Fig. 1d). Remarkably, F486V carried by BA.4/5 decreases hACE2 binding activity due to reduced hydrophobic interaction (Extended Data Fig. 1d). Potential hydrophilic interaction reduction could also be observed due to R493Q reversion. Notably, two recent reports claimed that BA.4/5 RBD and Spike (S2P) showed higher binding affinity to hACE2 compared to BA.1 and BA.2, due to L452R and R493Q reversion^18, 19^. Despite the discrepancy, it can be concluded that BA.2 subvariants and BA.4/5 were able to maintain high hACE2 binding affinity.

### NAb evasion of BA.2.12.1, BA.4 and BA.5

To probe the neutralization evasion ability of the recently emerged Omicron sublineages, we performed pseudovirus neutralization assays using D614G, BA.1, BA.1.1, BA.2, BA.3, BA.2.12.1, BA.2.13 and BA.4/BA.5 against the plasma obtained from 3-dose vaccinated individuals, BA.1 convalescents with previous vaccination, and vaccinated SARS convalescents (Supplementary Table 1). Plasma samples were collected 4 weeks after the booster shot or 4 weeks after COVID-19 hospital discharge. In individuals that received CoronaVac or ZF2001 booster six months after two doses of inactivated vaccine (CoronaVac), BA.1, BA.1.1 and BA.2 showed no significant difference in plasma neutralization resistance (Fig. 2a-b), concordant with previous reports^20, 21^. However, we found that BA.2 subvariants BA.2.13 and BA.2.12.1 showed increased immune evasion capability than BA.2, with BA.2.12.1 stronger than BA.2.13, and BA.4/BA.5 conferring even stronger antibody escape (Fig. 2a-b). The drop of neutralization titers is more obvious in the plasma obtained from individuals infected by BA.1 who had received 3-dose CoronaVac before infection (Fig. 2c), despite their significantly higher neutralization against D614G and BA.1 compared to the 3-dose vaccinees without BA.1 infection (Extended Data Fig. 2a). The plasma NT50 of BA.1 convalescents against BA.2.13, BA.2.12.1 and BA.4/5, compared to that against BA.1, was reduced by 2.0x, 3.7x and 8.0x fold, respectively. Interestingly, plasma from vaccinated SARS convalescents showed a different phenotype than normal vaccinees, such that BA.2 subvariants and BA.3/BA.4/BA.5 could cause a striking neutralization loss (Fig. 2d and Extended Data Fig. 2b). This suggests that certain mutations in BA.2 lineages and BA.3/4/5 may specifically evade broad sarbecovirus neutralizing antibodies, which are substantially enriched in vaccinated SARS convalescents^22^. Together, these observations indicate that the newly emerged BA.2.12.1 and BA.4/5 display stronger and distinct humoral immune evasion than BA.1.

**Fig. 2.**
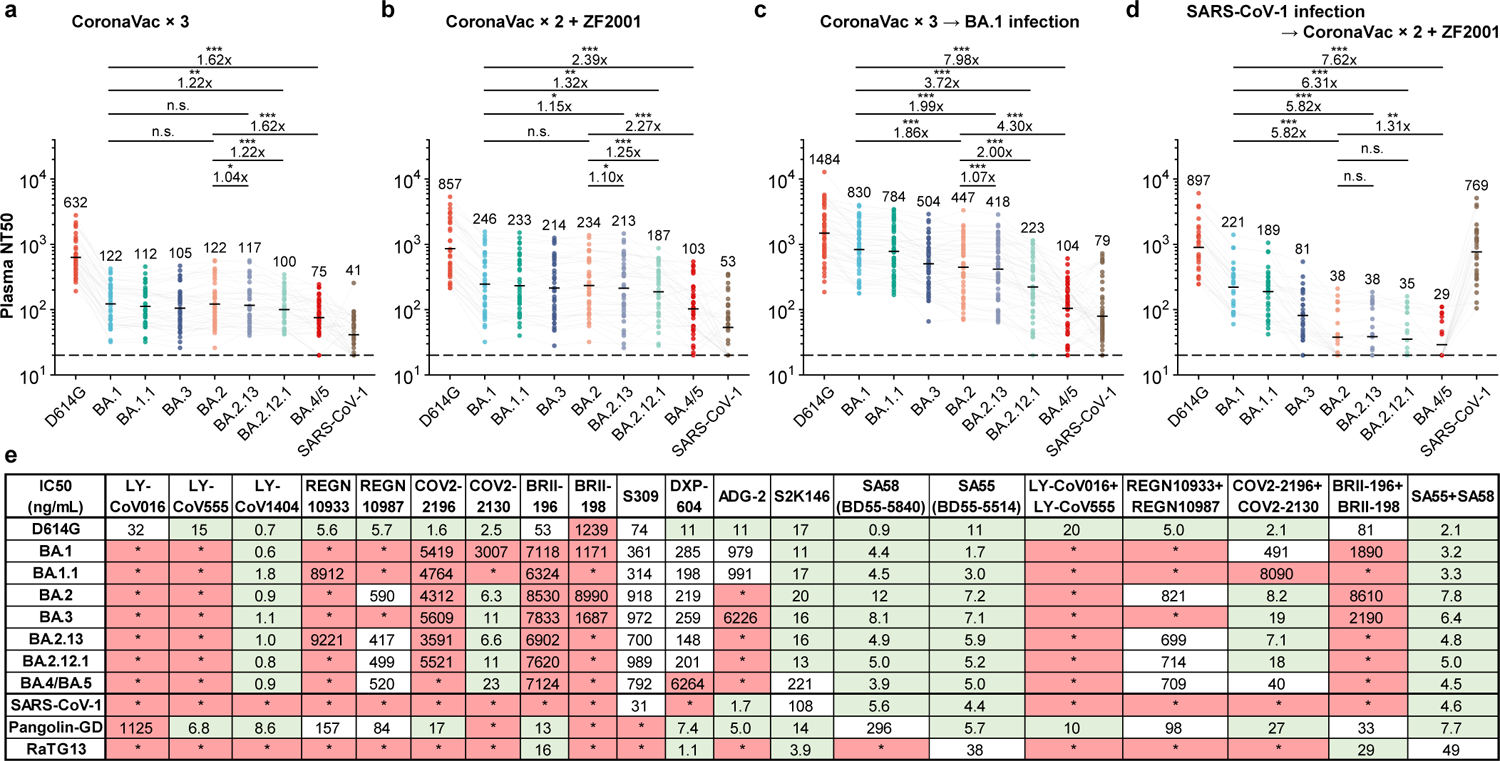
BA.2.12.1, BA.4 and BA.5 exhibit stronger antibody evasion than BA.2. **a-d**, Neutralizing titers against SARS-CoV-2 D614G, Omicron subvariants and SARS-CoV-1 pseudoviruses in plasma from vaccinated and convalescent individuals. **a**, Individuals who received 3 doses of CoronaVac (n=40). **b**, Individuals who received 2 doses of CoronaVac and 1 dose of ZF2001 (n=38). **c**, BA.1 convalescents who had received 3 doses of CoronaVac before infection (n=50). **d**, SARS convalescents who received 2 doses of CoronaVac and 1 dose of ZF2001 (n=28). P-values were calculated using two-tailed Wilcoxon signed-rank tests of paired samples. *, p<0.05; **, p<0.01; ***, p<0.001; n.s., not significant, p>0.05. Geometric mean titers (GMT) are labeled and annotated above each group of points. **e**, Neutralizing activity against SARS-CoV-2 variants and sarbecoviruses by therapeutic neutralizing antibodies; green, IC50 ≤ 30ng/mL; white, 30ng/mL < IC50 < 1,000ng/mL; red, IC50 ≥ 1,000ng/mL; *, IC50 ≥ 10,000ng/mL. All neutralization assays were conducted in biological duplicates.

Next, we examined the reaction difference in neutralizing activities of therapeutic antibodies against new Omicron subvariants (Fig. 2e). All of the seven Omicron subvariants displayed striking evasion against neutralization by Class 1 and 2 RBD antibodies, such that REGN-10933 (Casirivimab)^23^, LY-CoV016 (Etesevimab)^24^ and LY-CoV555 (Bamlanivimab)^25^, COV2-2196 (Tixagevimab)^5^ and BRII-196 (Amubarvimab)^26^ were strongly affected, while DXP-604^15, 27^ were evaded only by BA.4/5, showing reduced but still competitive efficacy against BA.1 and BA.2 subvariants. Two major antigenicity differences were observed between BA.1 and BA.2 subvariants. First, neutralizing antibodies targeting the linear epitope 440-449^3^, such as REGN-10987 (Imdevimab)^23^, COV2-2130 (Cilgavimab, component of Evusheld)^5^ and LY-CoV1404 (Bebtelovimab^4^) can neutralize BA.2 subvariants and BA.4/5. Second, BA.2 sublineages greatly reduce the efficacy of BA.1-effective broad sarbecovirus neutralizing antibodies, including ADG-2 (Adintrevimab)^28^ and S309 (Sotrovimab)^29^, except the ACE2-mimicking antibody S2K146^30^, which potently neutralize all BA.1 and BA.2 sublineages but showed reduced activity against BA.4/5, similar to DXP-604. BRII-196 and BRII-198 cocktail (Amubarvimab/Romlusevimab) were escaped by BA.2 sublineages^9^ and BA.3/BA.4/BA.5. Notably, LY-CoV1404^4^ demonstrated high potency against all assayed Omicron subvariants. In addition, our recently developed non-competing antibody cocktail isolated from vaccinated SARS convalescents, namely SA58 (BD55-5840, Class 3) and SA55 (BD55-5514, Class 1/4), displayed high potency against all Omicron subvariants and sarbecoviruses SARS-CoV-1, Pangolin-GD and RaTG13.

To delineate the underlying antibody evasion mechanism of BA.2.13, BA.2.12.1 and BA.4/BA.5, especially on how they escape the humoral immunity induced by BA.1 convalescents and vaccinated SARS convalescents, we started by isolating RBD-targeting NAbs from those individuals (Extended Data Fig. 3a)^27, 31^. First, antigen-specific memory B cells were isolated by fluorescence-activated cell sorting (FACS) from pooled PBMCs using double RBD^WT^+ selection for 3-dose vaccinees, RBD^WT^+ RBD^SARS^+ selection for vaccinated SARS convalescents and double RBD^BA.1^+ selection for BA.1 convalescents (Extended Data Fig. 3b). Secondly, we performed single-cell V(D)J sequencing (scVDJ-seq) with RBD^BA.1^ and RBD^WT^ feature barcodes to the CD27^+^/IgM^-^ antigen-specific memory B cells (Extended Data Fig. 3b). Thirdly, we extracted the productive heavy-light chain paired VDJ sequences and expressed the antibodies *in vitro* as human IgG1. Interestingly, during this process, we found that the majority of Omicron-reactive memory B cells from BA.1 convalescents who received 3-dose CornonaVac could also bind to WT RBD (Fig. 3a). In contrast, only around one-fourth of Omicron-reactive memory B cells isolated from unvaccinated BA.1 convalescents could bind to WT RBD (Fig. 3a). Also, the cross-reactive antigen-binding property could only be observed in IgM-CD27+ memory B cells, but not IgM+CD27-naïve B cells (Extended Data Fig. 2b). VDJ sequence analysis revealed significantly higher heavy chain V-domain somatic hypermutation (SHM) rates of BA.1/WT cross-reactive B cell receptors (BCRs) than that of BA.1-specific BCRs (Fig. 3b), which implies that cross-reactive memory B cells were further affinity-matured compared to BA.1-specific memory B cells. Together, these suggest that post-vaccination infection with Omicron BA.1 mainly recalls WT-induced memory B cells.

**Fig. 3.**
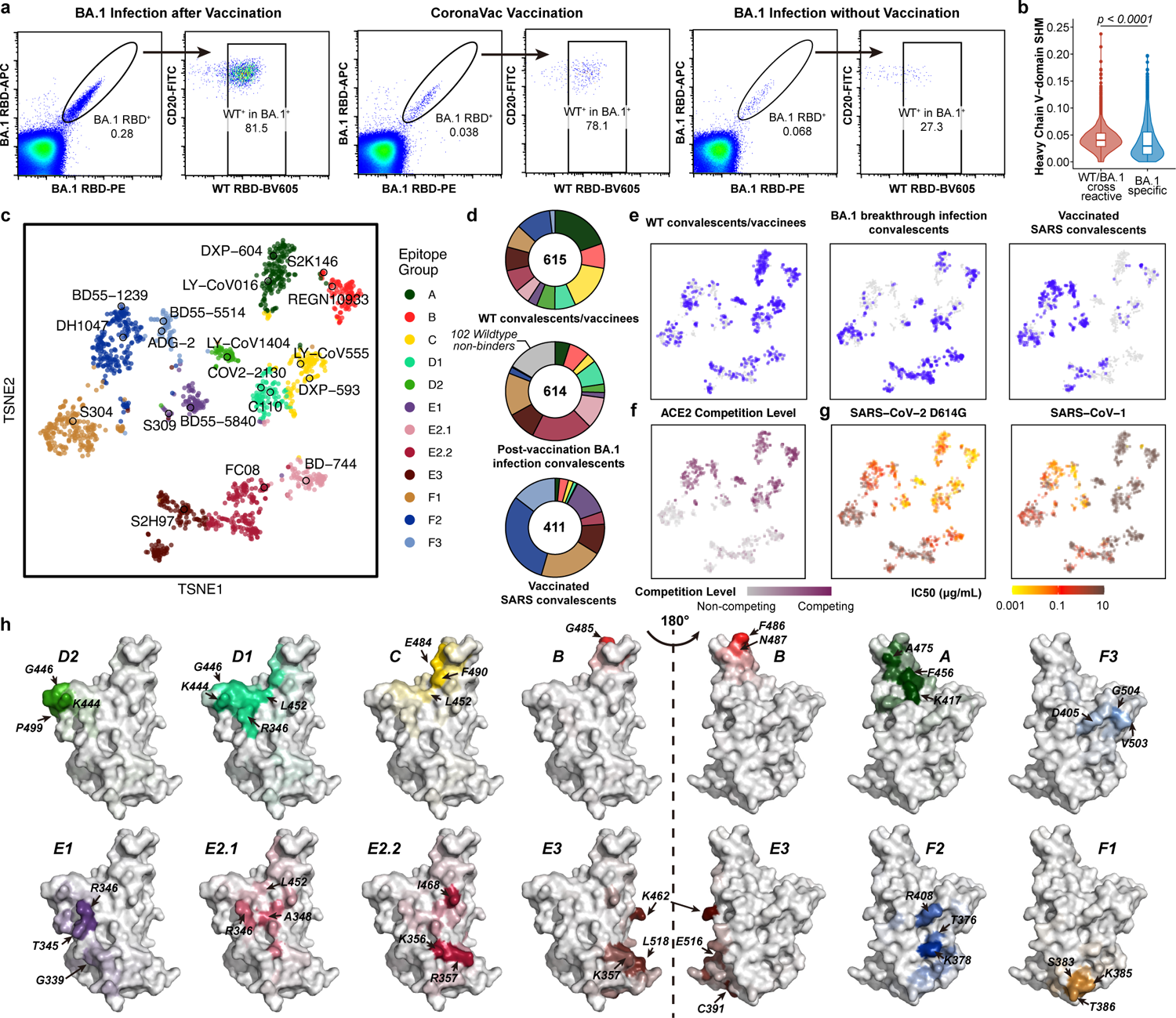
Isolation, characterization, and comprehensive epitope mapping of SARS-CoV-2 RBD antibodies. **a**, FACS analysis of pooled memory B cells (IgM^-^/CD27^+^) from BA.1 breakthrough infection convalescents, vaccinated individuals, and BA.1 convalescents without vaccination. **b**, Heavy chain V-domain somatic hypermutation (SHM) rate of BA.1-specific (n=968) and BA.1/WT cross-reactive (n=4782) BCRs obtained from 10X scVDJ-seq from post-vaccination BA.1 infection convalescents. P-value was calculated using two-tailed Wilcoxon rank-sum test. The 25 percentiles, medians and 75 percentiles are shown as boxplots. Kernel density estimation curves of the distribution are shown as violin plots. **c**, t-SNE and unsupervised clustering of SARS-CoV-2 wildtype RBD-binding antibodies. 12 epitope groups were identified based on deep mutational scanning of 1538 antibodies. **d-e**, Epitope distribution and projection of antibodies from wildtype convalescents, post-vaccination BA.1 infection convalescents, and vaccinated SARS convalescents. **f**, ACE2 competition level determined by competition ELISA (n=1286) were projected onto the t-SNE. **g**, Neutralizing activity against SARS-CoV-2 D614G (n=1509) and SARS-CoV-1 (HKU-39849, n=1457), respectively. **h**, Average mutational escape score projection of each epitope group on SARS-CoV-2 RBD (PDB: 6M0J). All neutralization assays were conducted in biological duplicates.

To further specify the epitope distribution of NAbs elicited by post-vaccination BA.1 infection, we applied high-throughput yeast-display-based deep mutational scanning (DMS) assays^2, 3^ and successfully determined the escaping mutation profiles of 1640 RBD-binding antibodies. Among these antibodies, 602 were from SARS-CoV-2 WT convalescents or 3-dose vaccinees, 614 from post-vaccination BA.1 convalescents, and 410 SARS/WT cross-reactive antibodies from vaccinated SARS convalescents (Supplementary Table 2). 14 antibodies with published DMS profiles are also included ^2, 32, 33^. It is important to note that, among the 614 antibodies from post-vaccination BA.1 convalescents, 102 are BA.1-specific and do not bind to WT RBD. The escaping mutation profiles of those 102 BA.1-specific NAbs were determined by DMS based on BA.1 RBD. The remaining 1538 RBD^WT^-reactive antibodies were unsupervised clustered into 12 epitope groups according to their WT-based mutational escaping profiles using t-distributed stochastic neighbor embedding (t-SNE) (Fig. 3c), adding 6 more epitope groups compared to our previous classification^3^.

Group A-C recapitulates our previous taxonomy^3^, in which the members mainly target the ACE2-binding motif^34–38^ (Fig. 3h). Group D antibodies, such as REGN10987, LY-CoV1404 and COV2-2130, bind to the linear epitope 440-449 on the RBD and are unsupervised expanded into D1 and D2 subgroups. Group D1 is more affected by R346 and L452, while D2 antibodies do not and interact more with P499 (Fig. 3h). Additionally, Group E and F are now expanded into E1-E3 and F1-F3, which covers the front and backside of RBD, roughly corresponding to Class 3 and Class 4, respectively^37^ (Fig. 3h). Group E1 corresponds to the S309 binding site, whose epitope involves G339, T345 and R346. Group E2 antibodies bind to the front chest of RBD^35^, where E2.1 is more affected by R346 and A348, while E2.2 is more affected by K356 and R357. Group E3 (S2H97 site) and F1 (S304 site) bind to highly conserved regions on the bottom of RBD, mainly contacting with K462/E516/L518, and S383/T385/K386, respectively. Group E1-E3 and F1 antibodies do not compete with ACE2 (Fig. 3f), while F2 and F3 antibodies are ACE2-competing and affected by T376, K378, D405, R408 and G504, corresponding to Class 1/4^39^. Pseuodovirus neutralizing efficacy of antibodies in each group against SARS-CoV-1, SARS-CoV-2 D614G, Pangolin-GD and RaTG13 is tested, and their binding capability to 22 sarbecovirus RBDs is measured through ELISA (Supplementary Table 2 and Supplementary Table 3). We found that antibodies of the same cluster have unified sarbecovirus neutralization potency and binding spectra (Fig. 3g and Extended Data Fig. 4). In total, five clusters of antibodies exhibiting broad sarbecovirus binding ability were identified, namely Groups E1, E3, F1, F2 and F3 (Extended Data Fig. 4), of which E1, F2 and F3 showed potent neutralizing activity against SARS-CoV-1 (Fig. 3g).

Importantly, we found that the individuals who experienced post-vaccination BA.1 infection displayed enrichment of Group E2.1, E2.2 and F1 antibodies (Fig. 3d-e), which do not compete with ACE2 (Fig. 3f). BA.1 does not harbor mutations on the epitopes of these NAb groups, which may explain why post-vaccination BA.1 infection is more likely to stimulate those NAbs. Though not enriched, the ACE2-competing Group B and D1 antibodies remain highly abundant. Since Group E2, D1 and B antibodies are sensitive to 452 and 486 mutations (Fig. 3h), it is highly possible that the newly emerged BA.2.12.1, BA.2.13, BA.4/BA.5 can specifically target those antibodies, rationalizing the huge loss in NT50 of BA.1 convalescents’ plasma against those variants (Fig. 2c).

To examine our hypothesis, we measured pseudovirus neutralization of those NAbs against BA.2.12.1, BA.2.13 and BA.4/BA.5, as well as the major Omicron variants BA.1, BA.1.1, BA.2 and BA.3 (Extended Data Fig. 5). Interestingly, NAbs from different epitope groups displayed distinct neutralizing activities against Omicron subvariants. Also, BA.1-stimulated antibodies (from BA.1 convalescents) and WT-stimulated (from WT convalescents or vaccinees, with or without previous SARS-CoV-1 infection) showed significantly higher potency and breadth in most epitope groups, confirming the higher affinity maturation (Extended Data Fig. 5).

Most WT-stimulated Group A, B and C NAbs were escaped by Omicron subvariants, while a subset showed broad Omicron effectiveness (Extended Data Fig. 5). Those broad NAbs are largely enriched by BA.1 stimulation and generally use similar heavy chain V genes compared to WT-stimulated antibodies and display higher convergence (Extended Data Fig. 6a-b). These broad ACE2-competing NAbs in Group A, B and C are also shown to be enriched in individuals who received a booster dose of mRNA vaccines^39^, which probably accounts for the high plasma neutralizing activity of 3-dose mRNA vaccinees against Omicron variants. Nevertheless, BA.1-stimulated groups B and C NAbs were significantly evaded by BA.4 due to F486V and L452R, concordant with results from DMS (Extended Data Fig. 7a-b), which explains the strong humoral immune evasion ability of BA.4/5.

Group D antibodies are most affected by G446S in BA.1, BA.1.1 and BA.3 (Fig. 4d); thus, these NAbs showed higher potency against BA.2 (Fig. 4a-b). However, D1 antibodies showed reduced efficacy against L452 substitutions, with L452M (BA.2.13) causing mild escapes, L452Q causing moderate escapes (BA.2.12.1), and L452R (BA.4/BA.5) causing severe escapes (Fig. 4c-d). In contrast, D2 antibodies, especially those stimulated by BA.1 infection, showed exceptional broad and potent neutralizing activity against all Omicron subvariants, such as LY-CoV1404 (Fig. 4b and Extended Data Fig. 5). Notably, although Group D2 NAbs displayed good breadth, their epitopes are not conserved among sarbecoviruses (Fig. 4d), similar to that of group D1, E2.1, and E2.2. This suggests that their exceptional breadth is possibly due to their rarity in WT and BA.1 convalescents (Fig. 3f), and these NAbs may be the next target for SARS-CoV-2 to escape by evolving specific mutations on their epitopes.

**Fig. 4.**
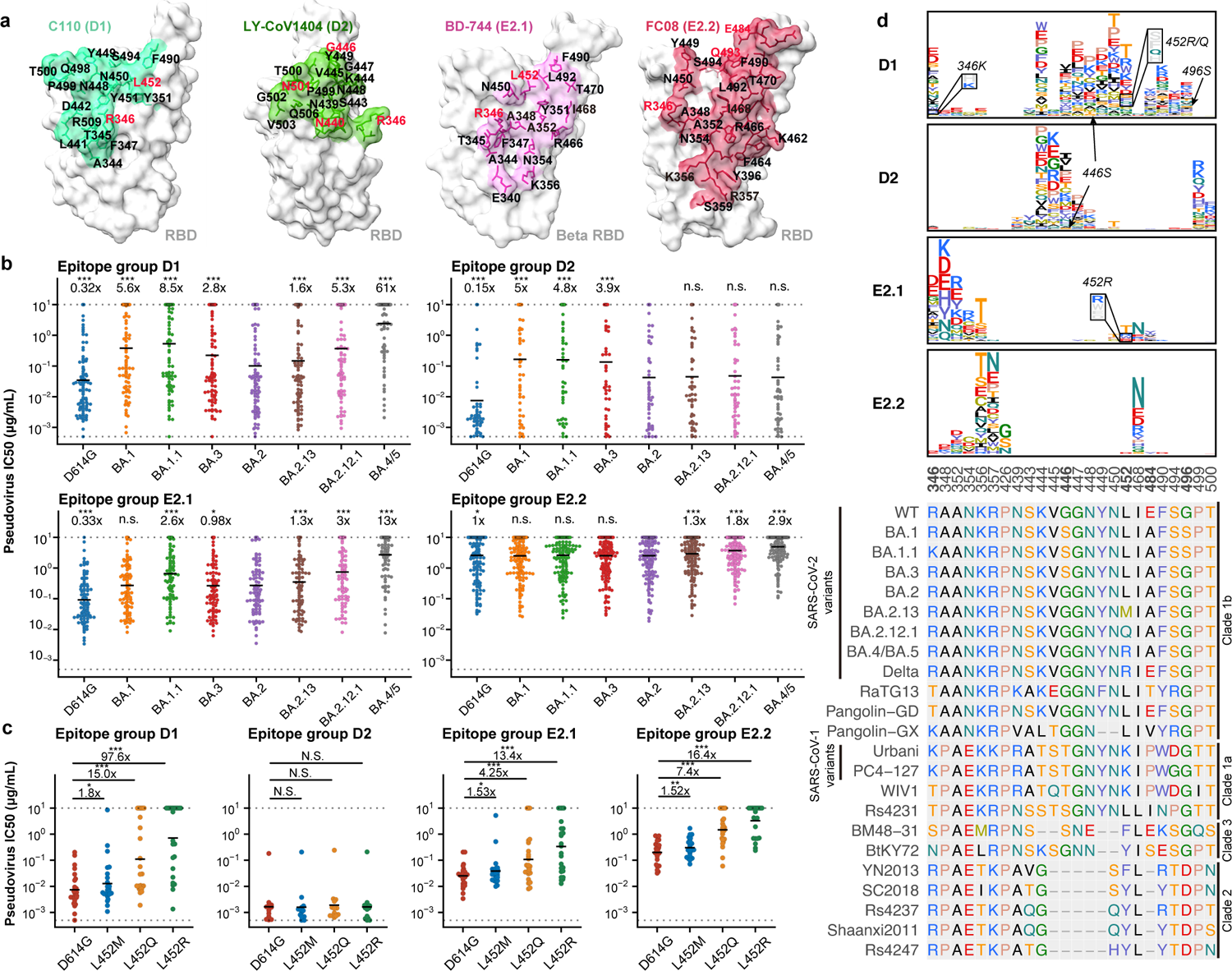
L452 mutants can evade cross-reactive NAbs elicited by BA.1 infection. **a**, Epitope of representative antibodies in group D1 (C110, PDB: 7K8V), D2 (LY-CoV1404, PDB: 7MMO), E2.1 (BD-744, PDB: 7EY0), and E2.2 (FC08, PDB: 7DX4). Residues highlighted in red indicate mutated sites in Omicron variants. **b**, Neutralizing activity of NAbs in group D1 (n=95), D2 (n=53), E2.1 (n=90) and E2.2 (n=161) against spike-pseudotyped SARS-CoV-2 variants. Geometric means of IC50 fold changes compared to BA.2 are annotated above the bars. P-values were calculated using a two-tailed Wilcoxon signed-rank test of paired samples, in comparison to IC50s against BA.2. **c**, Neutralizing activity of representative potent NAbs in group D1 (n=24), D2 (n=12), E2.1 (n=23) and E2.2 (n=23) against SARS-CoV-2 L452 mutants. Geometric mean of IC50 fold changes compared to IC50 against D614G are annotated above the points. P-values were calculated using a two-tailed Wilcoxon signed-rank test of paired samples. *, p<0.05; **, p<0.01; ***, p<0.001; n.s., not significant, p>0.05. **d**, Average escape maps at escape hotspots of antibodies in epitope group D1, D2, E2.1 and E2.2, and the corresponding MSA of various sarbecovirus RBDs. Height of each amino acid in the escape maps represents its mutation escape score. Mutated sites in Omicron variants are marked in bold. All neutralization assays were conducted in biological duplicates.

E2 antibodies bind to the chest of RBD^35^ (Fig. 4a), and their epitopes focus around R346, A348, A352, K356, R357 and I468 (Fig. 4d). Despite similar epitopes, E2.1 antibodies, especially those BA.1-stimulated, display significantly higher neutralizing potency than E2.2 (Fig. 4b). NAbs from the E2 Groups showed good breadth against SARS-COV-2 variants but not against BA.2.12.1 and BA.4/BA.5. L452 substitutions can cause large-scale escapes of E2.1 and E2.2 antibodies (Fig. 4c). Similar to the D1 epitope group, L452R and L452Q cause much stronger antibody evasion than L452M (Fig. 4c). Noteworthy, DMS does not reveal the L452 sensitivities of the E2.2 epitope group (Fig. 4d). Together, our results suggest that Omicron may have evolved mutations at L452 to specifically evade NAbs from D1 and E2, consequently maximizing Omicron BA.1 convalescents’ humoral immune evasion. Importantly, Group D1 and E2.1 antibodies also showed decreased efficacy against BA.1.1 compared to BA.1 (Fig. 4b), due to R346K, since both groups of NAbs are sensitive to the R346 substitution (Fig. 4a,d), which shed light on the prevalence of BA.1.1 after BA.1 in the United States.

### Omicron escapes broad sarbecovirus NAbs

In total, five clusters of antibodies were found to exhibit broad sarbecovirus neutralizing ability with diverse breadth, namely Group E1, E3, F1, F2 and F3 (Extended Data Fig. 4). While Group E3 and F1 antibodies demonstrated weak neutralizing activity against all variants due to their highly conserved binding sites (Extended Data Fig. 8a-c), we found that BA.1-effective E1, F2 and F3 NAbs, which are rare in WT and Omicron convalescents but enriched in vaccinated SARS convalescents, displayed a systematic reduction in neutralization activity against BA.2 subvariants and BA.3/BA.4/BA.5 (Fig. 2e and 5a-c). This observation explains the sharp NT50 drop of plasma from SARS convalescents against Omicron subvariants other than BA.1 (Fig. 2d). The mechanisms behind the neutralization loss of those broad sarbecovirus antibodies require investigation, which is critical for developing broad-spectrum sarbecovirus vaccines and antibody therapeutics.

**Fig. 5.**
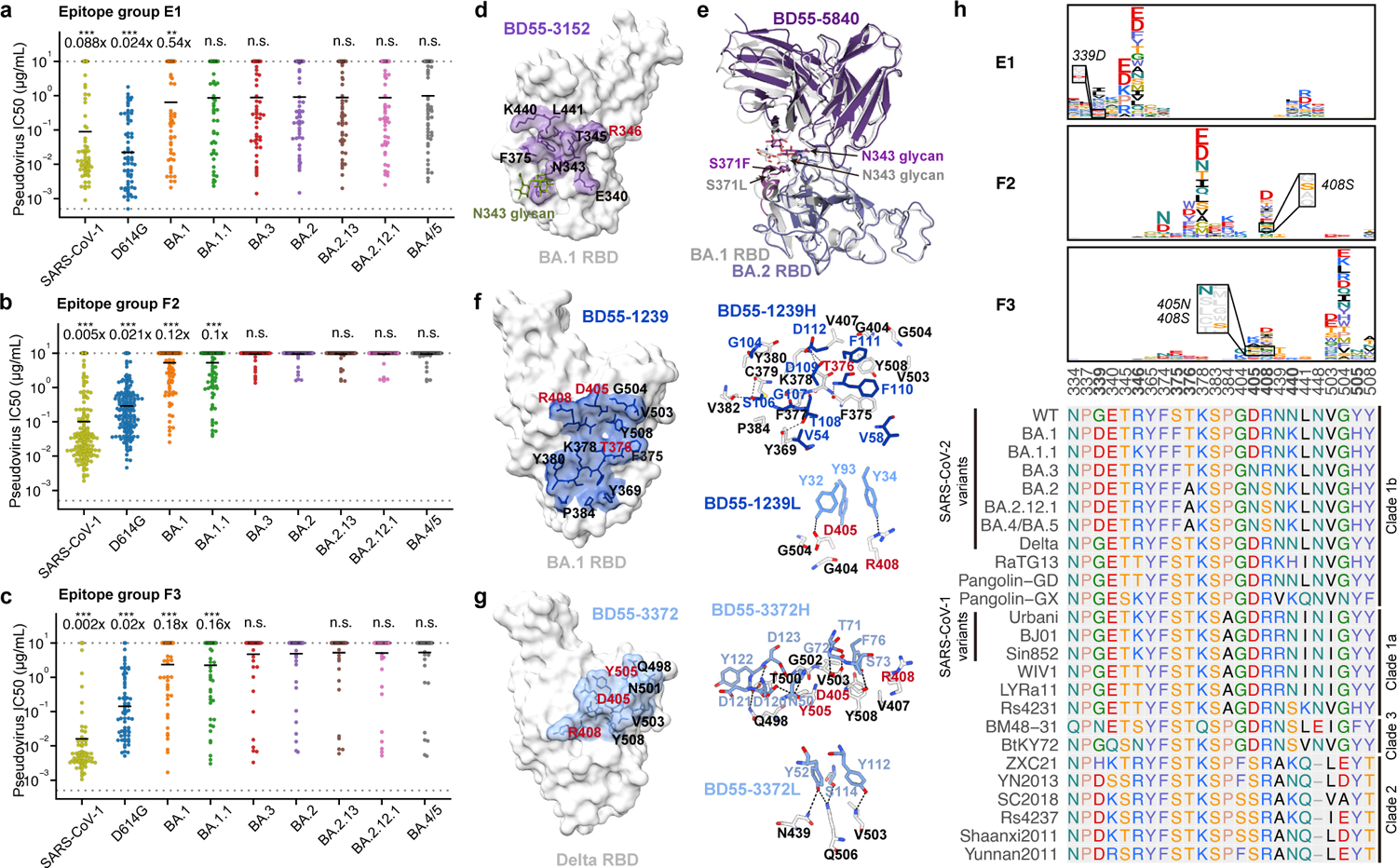
BA.2 subvariants can escape most broad sarbecovirus neutralizing antibodies. **a-c**, Neutralizing activity against SARS-CoV-1 and SARS-CoV-2 subvariants by NAbs in group E1 (**a**, n=70), F2 (**b**, n=171) and F3 (**c**, n=69). Geometric mean of IC50 fold changes compared to BA.2 are annotated above the bars. Geometric means of IC50 are labeled. P-values were calculated using a two-tailed Wilcoxon signed-rank test of paired samples, in comparison to IC50 against BA.2. *, p<0.05; **, p<0.01; ***, p<0.001; n.s., not significant, p>0.05. **d**, Epitope of Group E1 antibody BD55-3152 on SARS-CoV-2 BA.1 RBD. **e**, Structural overlay of BD55-5840 in complex of BA.1 and BA.2 RBD. **f-g**, Epitope and interactions on the binding interface of BD55-1239 (Group F2) and BD55-3372 (Group F3). Residues of the antibody are blue, and RBD residues are black or red. Residues highlighted in red indicate mutated sites in Omicron variants. **h**, Average escape maps of antibodies in epitope group E1, F2 and F3, and corresponding multiple sequence alignment (MSA) of various sarbecovirus RBDs. Height of each amino acid in the escape maps represents its mutation escape score. Mutated sites in Omicron subvariants are marked in bold. All neutralization assays were conducted in biological duplicates.

To study why BA.2 subvariants and BA.3/BA.4/BA.5 could systematically reduce the neutralization efficacy of E1 antibodies, we solved the cryo-EM structures of two BA.1 neutralizing E1 antibodies, BD55-3152 and BD55-5840 (SA58), in complex with BA.1 spike proteins using cryo-electron microscopy (cryo-EM) (Fig. 5d and Extended Data Fig. 9a-b). Like S309, E1 antibodies’ epitope involves an N-linked glycan on N343 (Fig. 5d). Besides, members of Group E1 are generally sensitive to the changes of G339, E340, T345 and especially R346, revealed by their escaping mutation profiles (Fig. 5h). Intriguingly, the newly acquired mutations of BA.2 do not overlap with the shared epitope of E1 antibodies, suggesting that the systematic reduction in neutralization is not caused by amino-acid substitution, but potentially due to structural alteration. To explore this hypothesis, we further determined the cryo-EM structure of the prefusion-stabilized BA.2 spike in complex with the BD55-5840 Fab (Fig. 5e). A structural comparison with the BA.1 RBD binding to BD55-5840 described above suggests that the 366-377 hairpin loop displays significant conformational differences due to S371F and T376A mutations (Fig. 5e and Extended Data Fig. 9d). The overall positions of residues 375-376 have been displaced by >3 Å, which likely further decreases the binding of F2/F3 NAbs in addition to the T376A side-chain substitution. As a result, the bulky Phe resulting from the S371F mutation interferes with the positioning of the glycan moiety attached to N343, which in turn budges the heavy chain of BD55-5840 upward. This may explain the reduction of the binding of BD55-5840 and S309, rationalizing their reduced neutralizing activity (Fig. 5a and Extended Data Fig. 9e). Importantly, the N343 glycan is critically recognized by almost all E1 neutralizing antibodies, including S309. Thus, this group of broad and potent neutralizing antibodies is likely affected by the S371F mutation in a systematic manner through N343-glycan displacement.

The epitopes of group F2 and F3 antibodies cover a continuous surface on the backside of RBD and can only bind to the up RBDs (Fig. 2b). To probe how BA.2 escapes antibodies of group F2 and F3, we solved the cryo-EM structure of two representative BA.1-effective antibodies in these groups, BD55-1239 from group F2, and BD55-3372 from group F3, in complex with the BA.1 and Delta spike protein respectively (Fig. 5f-g and Extended Data Fig. 9a). Group F2 antibodies can be escaped by RBD mutation involving T376, K378, and R408 (Fig. 5h). Indeed, these residues are all at the heart of BD55-1239’s epitope, and are fairly conserved across sarbecoviruses (Fig. 5h). Importantly, D405N and R408S harbored by Omicron BA.2 sublineages could alter the antigenic surface that disrupts the binding of F2 antibodies (Fig. 5f), hence completely abolishing the neutralizing capacity of F2 antibodies (Fig. 5b). Similarly, the D405N and R408S mutations harbored by BA.2 subvariants could interrupt the heavy-chain binding of F3 antibodies, causing large-scale escapes of BA.1-effective F3 neutralizing antibodies (Fig. 5c). The above observations were further validated by neutralizing activity against spike-pseudotyped VSV harboring D614G+D405N and D614G+R408S. As expected, Group E1 antibodies were affected by neither D405N nor R408S single substitution, while F2 and F3 antibodies displayed significantly decreased activity (Extended Data Fig. 9c). Nevertheless, several members of F3 antibodies are not sensitive to the D405N and R408S mutations of BA.2, making them good therapeutic drug candidates, such as BD55-5514 (SA55) (Fig. 2e). In sum, we revealed that S371F, D405N and R408S mutations harbored by BA.2 and emerging Omicron variants could induce large-scale escapes of broad sarbecovirus neutralizing antibodies, which is critical to the development of broad sarbecovirus antibody therapeutics and vaccines.

### BA.1-specific NAbs exhibit poor breadths

Besides the WT/BA.1 cross-reactive NAbs, it is also important to investigate the epitope distribution of BA.1-specific NAbs that do not react with WT RBD. To do so, we built the yeast display variants library based on RBD BA.1, and determined the escape mutation maps of 102 BA.1-specific antibodies. By integrating analysis of the whole dataset containing 1640 SARS-CoV-2 RBD antibodies, we got the embedded features of the BA.1-specific NAbs and performed clustering and t-SNE similarly (Fig. 6a). The 102 NAbs were clustered into four BA.1-specific epitope groups, named A^Omi^, B^Omi^, D^Omi^, and F3^Omi^, since these groups are highly related to their corresponding WT epitope groups (Fig. 6a and 6e). These antibodies are all ACE2-competing and display high BA.1 neutralization potency, but cannot neutralize D614G and SARS-CoV-1 (Fig. 6b-d), due to N417K/Y501N/H505Y for A^Omi^, A484E/K478T for B^Omi^, K440N for D^Omi^, and R498Q/Y501N for F3^Omi^, as indicated by average escape maps of each group (Fig. 6e-f). Although some of the previously circulating variants carry the mutations mentioned above, such as N501Y in Alpha, K417N/E484K/N501Y in Beta, and T478K in Delta, only a small subset of the antibodies exhibited neutralizing activity against them (Fig. 6e). Also, nearly all of the BA.1-specific NAbs showed poor cross-reactivity against other Omicron subvariants (Fig. 6d). Specifically, most antibodies in F3^Omi^/A^Omi^ are evaded by BA.2 subvariants and BA.3 possibly due to D405N, and antibodies in B^Omi^ are strongly escaped by BA.4 due to F486V. Some D^Omi^ antibodies might be affected by S446G and were evaded by BA.2 subvariants and BA.4, which were not captured by DMS (Fig. 6g). To further validate the results obtained by DMS, we constructed pseudoviruses based on BA.1 that carry reverting mutations of N417K, K440N, S446G, K478T, A484E, R498Q, Y501N and H505Y, as well as BA.1+D405N and BA.1+R408S. Unfortunately, BA.1+D405N could not generate high enough titers for further experiments despite multiple constructing attempts; thus, we constructed BA.2 + N405D instead. We found that the N417K, R498Q, Y501N and H505Y reversions indeed can lead to heavy evasion of most A^Omi^ and F3^Omi^ antibodies, consistent with DMS results (Fig. 6g). Also, K484E and K478T are the major escaping mutants responsible for the poor breadth of B^Omi^ NAbs (Fig. 6d). BA.1+S446G caused a small group of D^Omi^ antibodies to lose neutralization, while R498Q and K440N caused the majority of D^Omi^ NAbs not to bind to WT RBD. Importantly, BA.1+R408S did not show neutralization reduction to BA.1-specific NAbs, while BA.2+N405D could restore the neutralization potency of A^Omi^ and F3^Omi^ antibodies against BA.2, indicating that D405N is the main reason that caused their poor cross-reactivity among BA.2/BA.3/BA.4/BA.5 sublineages (Fig. 6d and 6g). Interestingly, these BA.1-specific NAbs displayed different heavy chain V gene usage compared to WT-reactive antibodies in the corresponding epitope group. Specifically, antibodies in A^Omi^ and B^Omi^ did not show significant convergence. IGHV3-53/3-66 only contributes to a small subset of A^Omi^ antibodies. Instead, D^Omi^ antibodies were dominated by IGHV2-70 and IGHV5-51, while IGHV4-59 for F^Omi^ (Extended Data Fig. 10). These three V genes also appeared in WT-reactive antibodies, but were relatively rare and did not show significant epitope enrichment (Extended Data Fig. 6a-b).

**Fig. 6.**
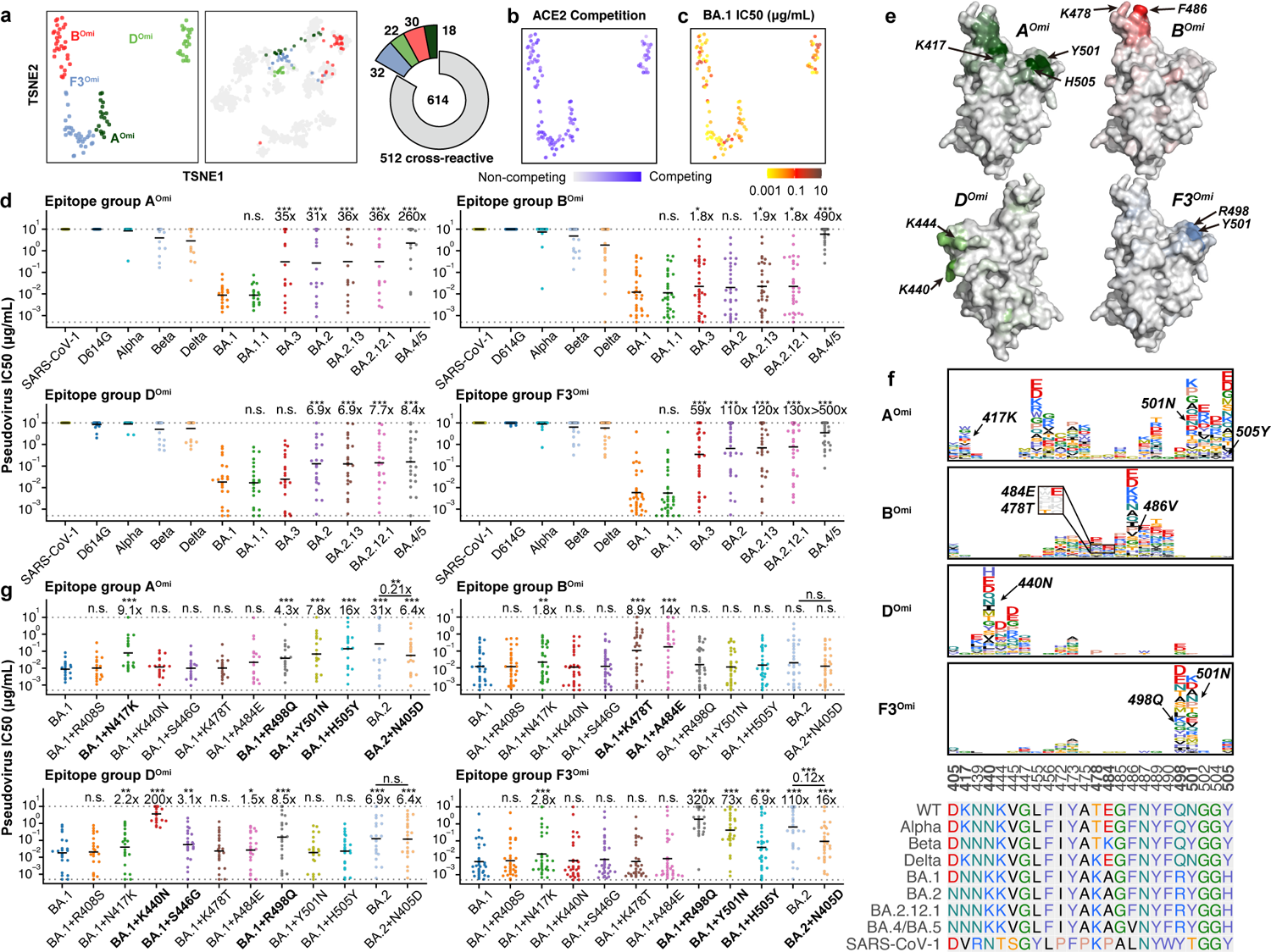
BA.1-specific antibodies elicited by BA.1 infection exhibit poor breadth. **a**, Four epitope groups were identified among 102 BA.1-specific NAbs, via k-means clustering and t-SNE of BA.1-RBD-based DMS profiles. **b-c**, Distribution of ACE2 competition level and neutralizing activities against BA.1. **d**, Pseudovirus neutralizing activities of BA.1-specific antibodies against SARS-CoV-1 and SARS-CoV-2 variants (A^Omi^, n=18; B^Omi^, n=30; D^Omi^, n=22; F3^Omi^, n=32). Geometric mean of IC50 fold changes compared to IC50 against BA.1 are annotated above the bars. Geometric means of IC50 are labeled. **e**, Average mutational escape score projection of each BA.1-specific epitope group on SARS-CoV-2 RBD (PDB: 7WPB). **f**, Averaged escape maps at escape hotspots of the 102 NAbs in four epitope groups, and corresponding MSA of various sarbecovirus RBDs. Height of each amino acid in the escape maps represents its mutation escape score. Mutated sites in Omicron variants are marked in bold. WT-related escaping mutations are highlighted. **g**, Neutralizing activities of BA.1-specific NAbs against BA.1 or BA.2 based pseudoviruses carrying single substitution (A^Omi^, n=18; B^Omi^, n=30; D^Omi^, n=22; F3^Omi^, n=32).. Geometric mean of IC50 fold changes compared to IC50 against BA.1 are annotated above the bars. All P-values were calculated using a two-tailed Wilcoxon signed-rank test of paired samples, in comparison with BA.1 IC50. *, p<0.05; **, p<0.01; ***, p<0.001; n.s., not significant, p>0.05. All neutralization assays were conducted in biological duplicates.

In this study, we showed that Omicron is continuously evolving under immune pressure, and rationalized the appearance of R346K (BA.1.1), L452 substitutions and F486V mutation, which all enabled stronger immune evasion. Unlike when Omicron first appeared, now Omicron sublineages could target the humoral immunity induced by Omicron itself, such as post-vaccination Omicron infection. The Omicron breakthrough infections mainly recall WT-induced memory B cells^40, 41^, which in turn narrows the diversity of antibodies elicited and may further facilitate the appearance of future mutants. These phenomena pose a great challenge to the currently established herd immunity through WT-based vaccination and BA.1/BA.2 infection. Similarly, these also suggest that Omicron BA.1-based vaccine may not be the ideal antigen for inducing broad-spectrum protection against emerging Omicron sublineages.

Here, by combining high-throughput single-cell sequencing and high-throughput yeast display-based deep mutational screening, we showcased the ability to decipher the complicated humoral immune repertoire elicited by Omicron infection and the underlying immune evasion mechanism of L452 and F486 mutations. The ability to dissect the entire humoral immunity into distinct antibody epitope groups greatly increases the resolution of antibody and mutational escape research. As we have shown, the antibodies in each epitope group show highly concordant attributes and features, largely facilitating the investigation of the immune evasion mechanism of circulating variants. The comprehensive data we provided in this research gives critical instructions to the development of broad-spectrum sarbecovirus vaccines and therapeutic antibodies.

## Supporting information

Supplementary Guide

## Methods

### Plasma and PBMC isolation

Blood samples were obtained from 40 volunteers who received 3 doses of CoronaVac, 39 individuals who received 2 doses of CoronaVac and 1 booster dose of ZF2001, 54 BA.1 convalescents who had received 3 doses of CoronaVac before BA.1 infection^42, 43^, and 30 SARS convalescents who received 2 doses of CoronaVac and 1 dose of ZF2001. The volunteers’ blood samples were obtained 4 weeks after the booster shot or 4 weeks after discharge from the hospital after BA.1 infection. COVID-19 disease severity was defined as asymptomatic, mild, moderate, severe and critical according to WHO living guidance for clinical management of COVID-19^44^. Relevant experiments regarding SARS convalescents and SARS-CoV-2 vaccinees were approved by the Beijing Ditan Hospital Capital Medical University (Ethics committee archiving No. LL-2021-024-02), the Tianjin Municipal Health Commission, and the Ethics Committee of Tianjin First Central Hospital (Ethics committee archiving No. 2022N045KY). Written informed consent was obtained from each participant in accordance with the Declaration of Helsinki. All participants provided written informed consent for the collection of information, storage and usage of their clinical samples for research purpose, and publication of data generated from this study.

Whole blood samples were mixed and subjected to Ficoll (Cytiva, 17-1440-03) gradient centrifugation after 1:1 dilution in PBS+2% FBS to isolate plasma and peripheral blood mononuclear cells (PBMC). After centrifugation, plasma was collected from upper layer and cells were harvested at the interface, respectively. PBMCs were further prepared through centrifugation, red blood cells lysis (InvitrogenTM eBioscienceTM 1X RBC Lysis Buffer, 00-4333-57) and washing steps. Samples were stored in FBS (Gibco) with 10% DMSO (Sigma) in liquid nitrogen if not used for downstream process immediately. Cryopreserved PBMCs were thawed in DPBS+2% FBS (Stemcell, 07905).

### Antibody isolation and recombinant production

SARS-CoV-1 and SARS-CoV-2 RBD cross-binding memory B cells were isolated from PBMC of SARS convalescents who received SARS-CoV-2 vaccine and BA.1 infected convalescents who had been vaccinated against COVID-19 prior to infection. Briefly, CD19+ B cells were isolated from PBMC with EasySep Human CD19 Positive Selection Kit II (STEMCELL, 17854). Every 10^6 B cells in 100 μL were then stained with 2.5 μL FITC anti-human CD19 antibody (BioLegend, 392508), 2.5 μL FITC anti-human CD20 antibody (BioLegend, 302304), 3.5 μL Brilliant Violet 421 anti-human CD27 antibody (BioLegend, 302824), 3 μL PE/Cyanine7 anti-human IgM antibody (BioLegend, 314532), 0.0052 μg biotinylated Ovalbumin (SinoBiological) conjugated with Brilliant Violet 605 Streptavidin (BioLegend, 405229), 0.0032 μg SARS-CoV-1 biotinylated RBD protein (His & AVI Tag) (SinoBiological, 40634-V27H-B) conjugated with PE-streptavidin (BioLegend, 405204), 0.0032 μg SARS-CoV-2 biotinylated RBD protein (His & AVI Tag) (SinoBiological, 40592-V27H-B) conjugated with APC-streptavidin (BioLegend, 405207), and 5 μL 7-AAD (Invitrogen, 00-6993-50). 7-AAD-, CD19/CD20+, CD27+, IgM-, OVA-, SARS-COV-1 RBD+, and SARS-CoV-2 RBD+ were sorted with MoFlo Astrios EQ Cell Sorter (Beckman Coulter).

SARS-CoV-2 BA.1 RBD binding memory B cells were isolated from BA.1 infected convalescents who received SARS-CoV-2. Briefly, CD19+ B cells were isolated with EasySep Human CD19 Positive Selection Kit II. Every 10^6 B cells in 100 μL solution were then stained with 3 μL FITC anti-human CD20 antibody (BioLegend, 302304), 3.5 μL Brilliant Violet 421 anti-human CD27 antibody (BioLegend, 302824), 2 μL PE/Cyanine7 anti-human IgM antibody (BioLegend, 314532), 2 μL PE/Cyanine7 anti-human IgD antibody(BioLegend, 348210), 0.0032 μg biotinylated SARS-CoV-2 BA.1 protein (His & AVI Tag) (SinoBiological, 40592-V49H7-B) conjugated with PE-streptavidin or APC-streptavidin (TotalSeq-C0971 Streptavidin, BioLegend, 405271 and TotalSeq-C0972 Streptavidin, BioLegend, 405273), 0.0032 μg SARS-CoV-2 WT biotinylated RBD protein (His & AVI Tag) conjugated with Brilliant Violet 605 Streptavidin and TotalSeq-C0973 Streptavidin (BioLegend, 405275) and TotalSeq-C0974 Streptavidin(BioLegend, 405277), 0.0052 μg biotinylated Ovalbumin conjugated with TotalSeq-C0975 Streptavidin (BioLegend, 405279) and 5 μL 7-AAD (Invitrogen, 00-6993-50). 7-AAD-, CD20+, CD27+, IgM-, IgD-, SARS-CoV-2 BA.1 RBD+ were sorted with MoFlo Astrios EQ Cell Sorter. FACS data were analyzed using FlowJo v10.8 (BD Biosciences).

Sorted B cells were then processed with Chromium Next GEM Single Cell V(D)J Reagent Kits v1.1 following the manufacturer’s user guide (10x Genomics, CG000208). Briefly, Cells sorted were resuspended in PBS after centrifugation. Gel beads-in-emulsion (GEMs) were obtained with 10X Chromium controller and then subjected to reverse transcription (RT). After GEM-RT clean up, RT products were subject to preamplification. After amplification and purification with SPRIselect Reagent Kit (Beckman Coulter, B23318) of RT products, B cell receptor (BCR) sequence (paired V(D)J) were enriched with 10X BCR primers. After library preparation, libraries were sequenced by Novaseq 6000 platform running Novaseq 6000 S4 Reagent Kit v1.5 300 cycles (Illumina, 20028312) or NovaSeq XP 4-Lane Kit v1.5 (Illumina, 20043131).

### B cell RNA and feature barcode data analysis

Using Cell Ranger (v6.1.1) pipeline, the mRNA fastq reads were processed and aligned to the human GRCh38 genome for gene expression profile. Genes expressed in less than 10 cells and cells expressed less than 100 genes or high-level mitochondria genes were removed, to filter out low-quality data. Raw counts were normalized and scaled with Seurat^45^ (v 4.0.3), while principal components analysis (PCA) and uniform manifold approximation and projection (UMAP) were performed for cluster and visualization. Cell types were identified using SingleR^46^ (v1.6.1) with Monaco human immune reference^47^. Feature barcode reads were also counted by Cell Ranger (v6.1.1) as antibody capture library, and a cell was considered to bind the corresponding antigen of dominant feature barcodes (>25% in this cell).

### Antibody sequence analysis

The antibody sequences obtained from 10X Genomics V(D)J sequencing were aligned to GRCh38 reference and assembled as immunoglobulin contigs by the Cell Ranger (v6.1.1) pipeline. Non-productive contigs and B cells that had multiple heavy chain or light chain contigs were filtered out of the analysis. V(D)J gene annotation was performed using NCBI IgBlast (v1.17.1) with the IMGT reference. Mutations on V(D)J nucleotide sequences were calculated by using the igpipeline, which compared the sequences to the closest germline genes and counted the number of different nucleotides. For antibodies from public sources whose original sequencing nucleotide sequences were not all accessible, the antibody amino acid sequences were annotated by IMGT/DomainGapAlign^48^ (v4.10.2) with default parameters. The V-J pairs were visualized by R package circlize (v0.4.10).

### Deep Mutational Scanning Library construction

Deep mutational scanning libraries were constructed as previously described^3^. Briefly, SARS-CoV-2 RBD mutant libraries were constructed from Wuhan-Hu-1 RBD sequence (GenBank: MN908947, residues N331-T531), and Omicron RBD mutant libraries were created in a similar way based on Wuhan-Hu-1 RBD sequence with the addition of G339D, S371L, S373P, S375F, K417N, N440K, G446S, S477N, T478K, E484A, Q493R, G496S, Q498R, N501Y, and Y505H mutations. Duplicated libraries were independently produced, theoretically containing 3819 possible amino acid mutations. Each RBD mutant was barcoded with a unique 26-nucleotide (N26) sequence and Pacbio sequencing was used to identify the correspondence of RBD mutant and N26 barcode. After mutant library transformation, ACE2 binders were enriched for downstream mutation profile experiment.

### High-throughput antibody-escape mutation profiling

The previously described high-throughput MACS (magnetic-activated cell sorting)-based antibody-escape mutation profiling system^3, 17^ was used to characterize mutation escape profile for neutralizing antibodies. Briefly, ACE2 binding mutants were induced overnight for RBD expression and washed followed by two rounds of Protein A antibody based negative selection and MYC-tag based positive selection to enrich RBD expressing cells. Protein A antibody conjugated products were prepared following the protocol of Dynabeads Protein A (Thermo Fisher, 10008D) and incubated with induced yeast libraries at room temperature for 30min with shaking. MYC-tag based positive selection was performed according to the manufacturer’s instructions (Thermo Fisher, 88843).

After three rounds of sequential cell sorting, the obtained cells were recovered overnight. Plasmids were extracted from pre- and post-sort yeast populations by 96-Well Plate Yeast Plasmid Preps Kit (Coolaber, PE053). The extracted plasmids were then used to amplify N26 barcode sequences by PCR. The final PCR products were purified with 1X AMPure XP magnetic beads (Beckman Coulter, A63882) and submitted to 75bp single-end sequencing at Illumina Nextseq 500 platform.

### Processing of deep mutational scanning data

Single-end Illumina sequencing reads were processed as previously described. Briefly, reads were trimmed into 16 or 26 bp and aligned to the reference barcode-variant dictionary with dms_variants package (v0.8.9). Escape scores of variants were calculated as F×(n_X,ab_ / N_ab_) / (n_X,ref_ / N_ref_), where n_X,ab_ and n_X,ref_ is the number of reads representing variant X, and N_ab_ and N_ref_ are the total number of valid reads in antibody-selected (ab) and reference (ref) library, respectively. F is a scale factor defined as the 99th percentiles of escape fraction ratios. Variants detected by less than 6 reads in the reference library were removed to avoid sampling noise. Variants containing mutations with ACE2 binding below −2.35 or RBD expression below −1 were removed as well, according to data previously reported. For RBD^BA.1^-based libraries, due to the lack of corresponding ACE2 binding and RBD expression data, we used the RBD expression of RBD^Beta^-based DMS as filter instead^49^, and did not perform the ACE2-binding filter. Mutations on residues that use different amino acids in Beta and BA.1 are not filtered, except R493P, S496P, R498P, H505P and all mutations on F375, which were excluded in the analysis due to low expression. Finally, global epistasis models were built using dms_variants package to estimate mutation escape scores. For most antibodies, at least two independent assays are conducted and single mutation escape scores are averaged across all experiments that pass quality control.

### Antibody clustering and visualization

Site total escape scores, defined as the sum of escape scores of all mutations at a particular site on RBD, were used to evaluate the impact of mutations on each site for each antibody. Each of these scores is considered as a feature of a certain antibody and used to construct a feature matrix **A**_N×M_ for downstream analysis, where N is the number of antibodies and M is the number of features (valid sites). Informative sites were selected using sklearn.feature_selection.VarianceThreshold of scikit-learn Python package (v0.24.2) with the variance threshold as 0.1. Then, the selected features were L2-normalized across antibodies using sklearn.preprocessing.normalize. The resulting matrix is referred as **A’**_N×M’_, where M’ is the number of selected features. The dissimilarity of two antibodies i, j is defined as 1-Corr(**A’**_i_,**A’**_j_), where Corr(**x**,**y**) is the Pearson’s correlation coefficient of vector **x** and **y**. We used sklearn.manifold.MDS to reduce the number of features from M’ to D=20 with multidimensional scaling under the above metric. Antibodies are clustered into 12 epitope groups using sklearn.cluster.KMeans of scikit-learn in the resulting D-dimensional feature space. Finally, these D-dimensional representations of antibodies were further embedded into two-dimensional space for visualization with t-SNE using sklearn.manifold.TSNE of scikit-learn. For the 102 BA.1-specific antibodies that were assayed with RBD^BA.1^-based yeast display library, the 20-dimensional embedding were generated using MDS with all 1640 antibodies’ DMS profile, but clustering and t-SNE were conducted independently. To project these antibodies onto the t-SNE space of 1538 antibodies assayed by RBD^WT^-based DMS, we calculated the pairwise Euclidean distance between 102 antibodies using RBD^BA.1^-based DMS and 1538 antibodies using RBD^WT^-based DMS in the 20-dimensional MDS space. The position of each BA.1-specific antibody in the original t-SNE space is defined as the average position of its ten nearest antibodies using RBD^WT^-based DMS. All t-SNE plots were generated by R package ggplot2 (v3.3.3).

### Pseudovirus neutralization assay

SARS-CoV-2 spike (GenBank: MN908947), Pangolin-GD spike (GISAID: EPI_ISL_410721), RaTG13 spike (GISAID: EPI_ISL_402131), SARS-CoV-1 spike (GenBank: AY278491), Omicron BA.1 spike (A67V, H69del, V70del, T95I, G142D, V143del, Y144del, Y145del, N211del, L212I, ins214EPE, G339D, S371L, S373P, S375F, K417N, N440K, G446S, S477N, T478K, E484A, Q493R, G496S, Q498R, N501Y, Y505H, T547K, D614G, H655Y, N679K, P681H, N764K, D796Y, N856K, Q954H, N969K, L981F), BA.2 spike (GISAID: EPI_ISL_7580387, T19I, L24S, del25-27, G142D, V213G, G339D, S371F, S373P, S375F, T376A, D405N, R408S, K417N, N440K, G446S, S477N, T478K, E484A, Q493R, Q498R, N501Y, Y505H, D614G, H655Y, N679K, P681H, N764K, D796Y, Q954H, N969K), BA.1.1 spike (BA.1+R346K), BA.3 spike (A67V, del69-70, T95I, G142D, V143del, Y144del, Y145del, N211del, L212I, G339D, S371F, S373P, S375F, D405N, K417N, N440K, G446S, S477N, T478K, E484A, Q493R, Q498R, N501Y, Y505H, D614G, H655Y, N679K, P681H, N764K, D796Y, Q954H, N969K), BA.2.12.1 spike (BA.2+L452Q+S704L), BA.2.13 spike (BA.2+L452M), BA.4 spike (T19I, L24S, del25-27, del69-70, G142D, V213G, G339D, S371F, S373P, S375F, T376A, D405N, R408S, K417N, N440K, G446S, L452R, S477N, T478K, E484A, F486V, Q498R, N501Y, Y505H, D614G, H655Y, N679K, P681H, N764K, D796Y, Q954H, N969K) plasmid is constructed into pcDNA3.1 vector. G*ΔG-VSV virus (VSV G pseudotyped virus, Kerafast) is used to infect 293T cells (American Type Culture Collection [ATCC], CRL-3216), and spike protein expressing plasmid was used for transfection at the same time. After culture, the supernatant containing pseudovirus was harvested, filtered, aliquoted, and frozen at −80°C for further use.

Pseudovirus detection of Pangolin-GD and RaTG13 was performed in 293T cells overexpressing human angiotensin-converting enzyme 2 (293T-hACE2 cells). Other pseudovirus neutralization assays were performed using the Huh-7 cell line (Japanese Collection of Research Bioresources [JCRB], 0403).

Monoclonal antibodies or plasma were serially diluted (5-fold or 3-fold) in DMEM (Hyclone, SH30243.01) and mixed with pseudovirus in 96-well plates. After incubation at 5% CO_2_ and 37℃ for 1 h, digested Huh-7 cell (Japanese Collection of Research Bioresources [JCRB], 0403) or 293T-hACE2 cells (AmericanTypeCultureCollection[ATCC],CRL-3216) were seeded. After 24 hours of culture, supernatant was discarded and D-luciferin reagent (PerkinElmer, 6066769) was added to react in the dark, and the luminescence value was detected using a microplate spectrophotometer (PerkinElmer, HH3400). IC50 was determined by a four-parameter logistic regression model using PRISM (versions 9.0.1).

### ELISA

To detect the broad-spectrum binding of the antibodies among Sarbecovirus, we entrusted SinoBiological Technology Co., Ltd. to synthesize a panel of 20 sarbecovirus RBDs (Supplementary Table 3). According to the sequence of 20 RBDs, a set of nested primers was designed. The coding sequences were obtained by the overlap-PCR with a 6xHis tag sequence to facilitate protein purification. The purified PCR products were ligated to the secretory expression vector pCMV3 with CMV promoter, and then transformed into E. coli competent cells XL1-blue. Monoclones with correct transformation were cultured and expanded, and plasmids were extracted. Healthy HEK293F cells were passaged into a new cell culture and grown in suspension at 37 °C, 120 RPM, 8% CO2 to logarithmic growth phase and transfected with the recombinant constructs by using liposomal vesicles as DNA carrier. After transfection, the cell cultures were followed to assess the kinetics of cell growth and viability for 7 days. The cell expression supernatant was collected, and after centrifugation, passed through a Ni column for affinity purification. The molecular size and purity of eluted protein was confirmed by SDS-PAGE. Production lot numbers and concentration information of the 20 Sarbecovirus proteins are shown in Supplementary Table 4. The WT RBD in the article is SARS-CoV-2 (2019-nCoV) Spike RBD-His Recombinant Protein (SinoBiological, 40592-V08H).

A panel of 21 sarbecovirus RBDs (Supplementary Table 3) in PBS was pre-coated onto ELISA Plates (NEST, 514201) at 4℃ overnight. The plates were washed and blocked. Then 1μg/ml purified antibodies or serially diluted antibodies were added and incubated at room temperature (RT) for 20min. Next, Peroxidase-conjugated AffiniPure Goat Anti-Human IgG (H+L) (JACKSON, 109-035-003) was applied and incubated at RT for 15min. Tetramethylbenzidine (TMB) (Solarbio, 54827-17-7) was added onto the plates. The reaction was terminated by 2 M H_2_SO_4_ after 10min incubation. Absorbance was measured at 450 nm using Ensight Multimode Plate Reader (PerkinElmer, HH3400). ELISA OD450 measurements at different antibody concentration for a particular antibody-antigen pair are fit to the model y=Ac^n^/(c^n^ + E^n^) using R package mosaic (v1.8.3), where y is OD450 values and c is corresponding antibody concentration. A, E, n are parameters, where E is the desired EC_50_ value for the specific antibody and antigen.

### Antibody-ACE2 competition for RBD

Omicron-RBD (Sino Biological, 40592-V08H121) protein in PBS was immobilized on the ELISA plates at 4℃ overnight. The coating solution was removed and washed three times by PBST and the plates were then blocked for 2 h. After blocking, the plates were washed five times, and the mixture of ACE2-biotin (Sino Biological, 10108-H27B-B) and serially diluted competitor antibodies was added followed by 30min incubation at RT. Then Peroxidase-conjugated Streptavidin (Jackson ImmunoResearch, 016-030-084) was added into each well for another 20min incubation at RT. After washing the plates for five times, Tetramethylbenzidine (TMB) (Solarbio, 54827-17-7) was added into each well. After 10 min, the reaction was terminated by 2M H_2_SO_4_. Absorbance was measured at 450 nm using Ensight Multimode Plate Reader (PerkinElmer, HH3400). The ACE2 competition coefficient is calculated as (B-A)/B, where B is the OD450 value under 0.3ug/ml antibody concentration and A is the OD450 value under 6ug/ml antibody concentration.

### Biolayer Interferometry

Biolayer interferometry assays were performed on Octet® RED 384 Protein Analysis System (Fortebio) according to the manufacturer’s instruction. To measure the binding affinities, monoclonal antibodies were immobilized onto Protein A biosensors (Fortebio) and the fourfold serial dilutions of Omicron S trimer (BA.1 and BA.2) in PBS were used as analytes. Data were collected with Octet Acquisition 9.0 (Fortebio) and analyzed by Octet Analysis 9.0 (Fortebio) and Octet Analysis Studio 12.2 (Fortebio).

### S trimer thermal stability assay

The thermal stability assay was performed to detect the exposed hydrophobic residues by an MX3005 qPCR instrument (Agilent, Santa Clara, USA) with SYPRO Red (Invitrogen, Carlsbad, USA) as fluorescent probes. Here, we set up 25 μL reaction system (pH=8.0) which contained 5 μg of target protein (S trimer of Omicron lineage), 1000x SYPRO Red, and ramped up the temperature from 25°C to 99°C. Fluorescence was recorded in triplicate at an interval of 1°C.

### Surface plasmon resonance

Human ACE2 was immobilized onto CM5 sensor chips using a Biacore 8K (GE Healthcare). Serial dilutions of purified S trimer or RBD of Omicron lineages were injected, ranging in concentrations from 100 to 6.25 nM. The response units were recorded at room temperature using BIAcore 8K Evaluation Software (v3.0.12.15655; GE Healthcare), and the resulting data were fitted to a 1:1 binding model using BIAcore 8K Evaluation Software (v3.0.12.15655; GE Healthcare).

### Protein expression and purification for cryo-EM study

The S6P expression construct encoding the SARS-CoV-2 spike ectodomain (residues 1-1208) with six stabilizing Pro substitutions (F817P, A892P, A899P, A942P, K986P, and V987P) and a “GSAS” substitution for the furin cleavage site (residues 682–685) was previously described^15^. The Delta specific mutations (T19R, G142D, 156del, 157del, R158G, L452R, T478K, D614G, P681R, D950N) were introduced into this construct using site-directed mutagenesis. The S6P expression construct containing the Omicron BA.1 mutations (A67V, H69del, V70del, T95I, G142D, V143del, Y144del, Y145del, N211del, L212I, ins214EPE, G339D, S371L, S373P, S375F, K417N, N440K, G446S, S477N, T478K, E484A, Q493R, G496S, Q498R, N501Y, Y505H, T547K, D614G, H655Y, N679K, P681H, N764K, D796Y, N856K, Q954H, N969K, L981F) were assembled from three synthesized DNA fragments. The S6P expression construct containing the Omicron BA.2 mutations (T19I, L24S, del25-27, G142D, V213G, G339D, S371F, S373P, S375F, T376A, D405N, R408S, K417N, N440K, G446S, S477N, T478K, E484A, Q493R, Q498R, N501Y, Y505H, D614G, H655Y, N679K, P681H, N764K, D796Y, Q954H, N969K) were assembled from three synthesized DNA fragments. The S6P expression construct containing the Omicron BA.4/5 mutations (T19I, L24S, del25-27, del69-70, G142D, V213G, G339D, S371F, S373P, S375F, T376A, D405N, R408S, K417N, N440K, G446S, L452R, S477N, T478K, E484A, F486V, Q498R, N501Y, Y505H, D614G, H655Y, N658S, N679K, P681H, N764K, D796Y, Q954H, N969K) were assembled from three synthesized DNA fragments. The expression constructs encoding the SARS-CoV spike ectodomain (residues 1-1195)^50^ was kindly provided by Prof. X. Wang (Tsinghua University), and two stabilizing Pro substitutions (K968P, V969P) was engineered into this construct using mutagenesis. For protein production, these expression plasmids, as well as the plasmids encoding the antigen-binding fragments (Fabs) of the antibodies described in this paper, were transfected into the HEK293F cells using polyethylenimine (Polysciences). The conditioned media were harvested and concentrated using a Hydrosart ultrafilter (Sartorius), and exchanged into the binding buffer (25 mM Tris, pH 8.0, and 200 mM NaCl). Protein purifications were performed using the Ni-NTA affinity method, followed by gel filtration chromatographies using either a Superose 6 increase column (for the spike proteins) or a Superose 200 increase column (for the Fabs). The final buffer used for all proteins is 20 mM HEPES, pH 7.2, and 150 mM NaCl.

### Cryo-EM data collection, processing, and structure building

The samples for cryo-EM study were prepared essentially as previously described^15, 51^ (Supplementary Table 4). All EM grids were evacuated for 2 min and glow-discharged for 30 s using a plasma cleaner (Harrick PDC-32G-2). Four microliters of spike protein (0.8 mg/mL) was mixed with the same volume of Fabs (1 mg/mL each), and the mixture was immediately applied to glow-discharged holy-carbon gold grids (Quantifoil, R1.2/1.3) in an FEI Vitrobot IV (4 °C and 100% humidity). Data collection was performed using either a Titan Krios G3 equipped with a K3 direct detection camera, or a Titan Krios G2 with a K2 camera, both operating at 300 kV. Data processing was carried out using cryoSPARC (v3.2.1)^52^. After 2D classification, particles with good qualities were selected for global 3D reconstruction and then subjected to homogeneous refinement. To improve the density surrounding the RBD-Fab region, UCSF Chimera (v1.16)^53^ and Relion (v3.1)^54^ were used to generate the masks, and local refinement was then performed using cryoSPARC (v3.2.1). Coot (v0.8.9.2)^55^ and Phenix (v1.20)^56^ were used for structural modeling and refinement. Figures were prepared using USCF ChimeraX (v1.3)^57^ and Pymol (v2.6.0a0, Schrödinger, LLC.).

### Molecular dynamics simulation

Models of the RBD from BA.1, BA.2, BA.3, BA.2.13, BA.2.12.1 and BA.4 in complex with ACE2 were firstly referred to the cryo-EM structure of BA.1-hACE2 (PDB code: 7WGB) and then checked by WHAT IF Web Interface to remove atomic clashes. After that, the structures were simulated by the software GROMACS-2021^58^. Briefly, OPLS force field with TIP3P water model was selected to prepare the dynamic system. After that Na+ and Cl^-^ ions were added into the system to make the system electrically neutralized. Then, energy minimization using the steepest descent algorithm was carried out until the maximum force of 1,000 kJ mol^-1^ has been achieved. NVT ensemb1e via the Nose-Hoover method at 300 K and NPT ensemble at 1 bar with the Parinello-Rahman algorithm were employed successively to make the temperature and the pressure equilibrated, respectively. Finally, MD production runs of 10 ns were performed with random initial velocities and periodic boundary conditions. The non-bonded interactions were treated using Verlet cut-off scheme, while the long-range electrostatic interactions were treated using particle mesh Ewald (PME) method^58^. The short-range electrostatic and van der Waals interactions were calculated with a cut-off of 12 Å. All the 6 models were simulated in the same protocol.

## Data availability

Processed mutation escape scores can be downloaded at https://github.com/jianfcpku/SARS-CoV-2-RBD-DMS-broad. Raw Illumina and PacBio sequencing data are available on NCBI Sequence Read Archive BioProject PRJNA804413. We used vdj_GRCh38_alts_ensembl-5.0.0 as the reference of V(D)J alignment, which can be obtained from https://support.10xgenomics.com/single-cell-vdj/software/downloads/latest. IMGT/DomainGapAlign is based on the built-in lastest IMGT antibody database, and we let the “Species” parameter as “Homo sapiens” while kept the others as default. Public deep mutational scanning datasets involved in the study from literature could be downloaded at https://media.githubusercontent.com/media/jbloomlab/SARS2_RBD_Ab_escape_maps/main/processed_data/escape_data.csv. Public structures involved in this manuscript were downloaded from Protein Data Bank with accession codes 6M0J, 7K8V, 7MMO, 7EY0, 7DX4, 7M7W, 7JW0, 7WPB, 7WGB. Cryo-EM density maps have been deposited in the Electron Microscopy Data Bank with accession codes EMD-33210, EMD-33211, EMD-33212, EMD-33213, EMD-33323, EMD-33324, EMD-33325, EMD-32732, EMD-32738, EMD-32734, EMD-32718, and EMD-33019, respectively. Structural coordinates have been deposited in the Protein Data Bank with accession codes 7XIW, 7XIX, 7XIY, 7XIZ, 7XNQ, 7XNR, 7XNS 7WRL, 7WRZ, 7WRO, 7WR8 and 7X6A.

## Code availability

Python and R scripts for analyzing escaping mutation profile data and reproducing figures in this manuscript are available at https://github.com/jianfcpku/SARS-CoV-2-RBD-DMS-broad.

## Ethical Statement

This study was approved by the Ethics Committee of Beijing Ditan Hospital affiliated to Capital Medical University (Ethics committee archiving No. LL-2021-024-02), the Tianjin Municipal Health Commission, and the Ethics Committee of Tianjin First Central Hospital (Ethics committee archiving No. 2022N045KY). Informed consent was obtained from all human research participants.

## Acknowledgments

We thank J. Bloom for his gift of the yeast SARS-CoV-2 WT RBD libraries. We thank Sino Biological for the technical assistance on mAbs and RBD expression. We thank J. Luo and H. Lv for the help with flow cytometry. This project is financially supported by the Ministry of Science and Technology of China (CPL-1233).

## Author contributions

Y.Cao and X.S.X designed the study. Y.Cao, F.J. and X.S.X wrote the manuscript with inputs from all authors. Y.Cao. and F.S. coordinated the expression and characterization of the neutralizing antibodies. J.W. (BIOPIC), F.J., L.Z., H.S. performed and analyzed the yeast display screening experiments. Y.Y., T.X., P.W., J.W. (Changping Laboratory), R.A., Y.W., J.Z., N.Z., R.W., X.N., L.Y., C.L., X.S. L.Z., F.S. performed the neutralizing antibody expression and characterization, including pseudovirus neutralization and ELISA. Y.Y., W.H., Q.L., Y.W. prepared the VSV-based SARS-CoV-2 pseudovirus. A.Y., Y.W., S.Y., R.A., W.S. performed and analyzed the antigen-specific single B cell VDJ sequencing. S.D., P.L., Z.Z., L.W., R.F., Z.L., X.W., J.X. performed the structural analyses. X.H., W.Z., D.Z., and R.J. recruited the SARS convalescents and SARS-CoV-2 vaccinees. X.C. and Z.S. recruited the Omicron BA.1 convalescents. X.C., Y.Chai, Y.H., and Y.S. isolated PBMC from BA.1 convalescents. Q.G. proofed the manuscript.

## Extended Data Figures

**Extended Data Fig. 1.**
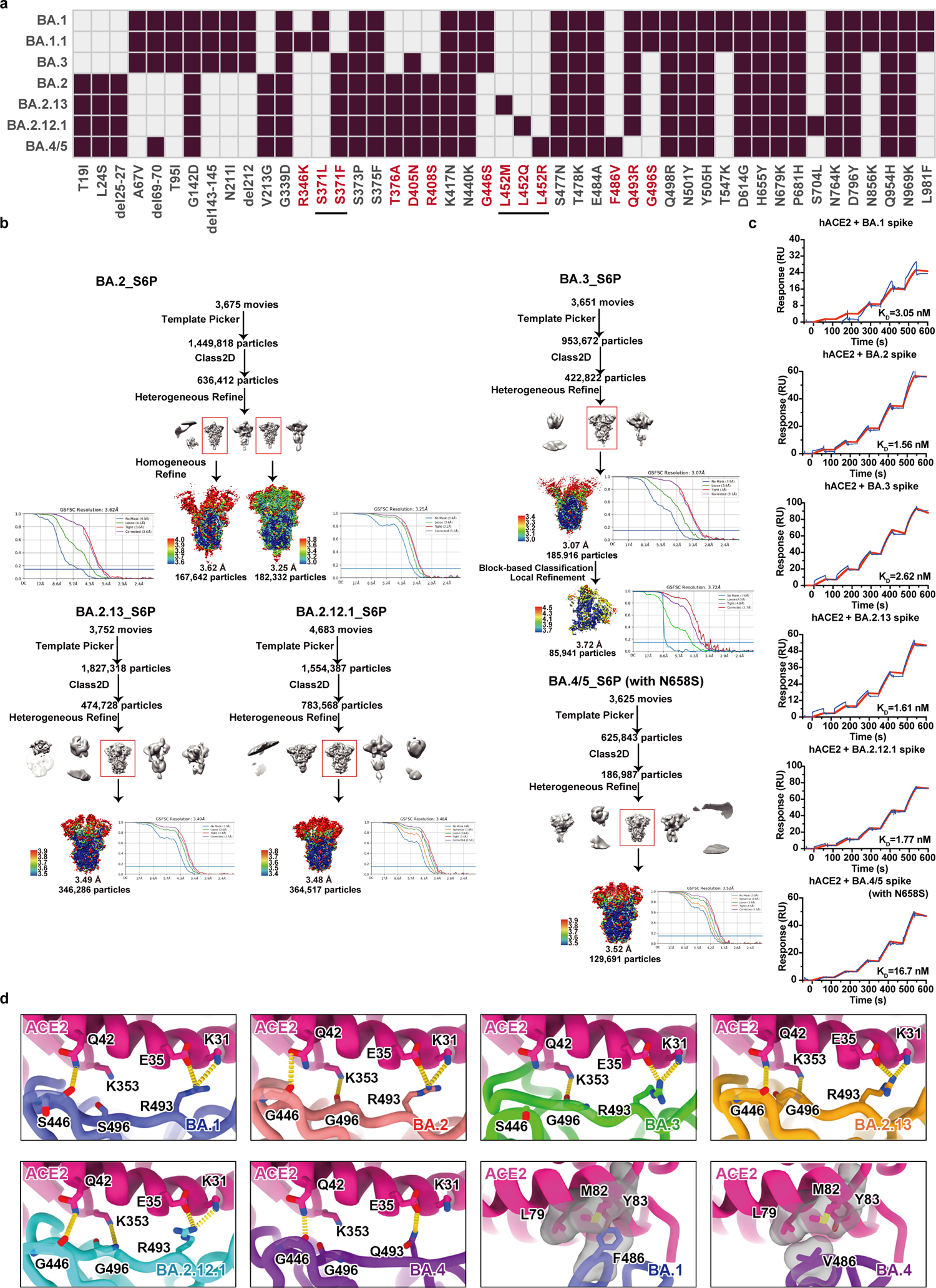
Structures and ACE2 binding of emerging Omicron subvariants spike glycoprotein. **a**, Mutations on the spike glycoprotein of SARS-CoV-2 Omicron subvariants. Residues that are not identical among Omicron subvariants are colored red. **b**, Workflow to generate cryo-EM structure of BA.2, BA.3, BA.2.13, BA.2.12.1, BA.4/5 spike glycoprotein trimer with S6P and R683A, R685A substitutions. **c**, Binding affinities of Omicron variants spike trimers to hACE2 measured by SPR. SPR analyses were conducted in biological duplicates. **d**, MD simulated interactions between hACE2 and RBD of Omicron variants. Structures of the RBD from Omicron variants and hACE2 are shown as ribbons.

**Extended Data Fig. 2.**
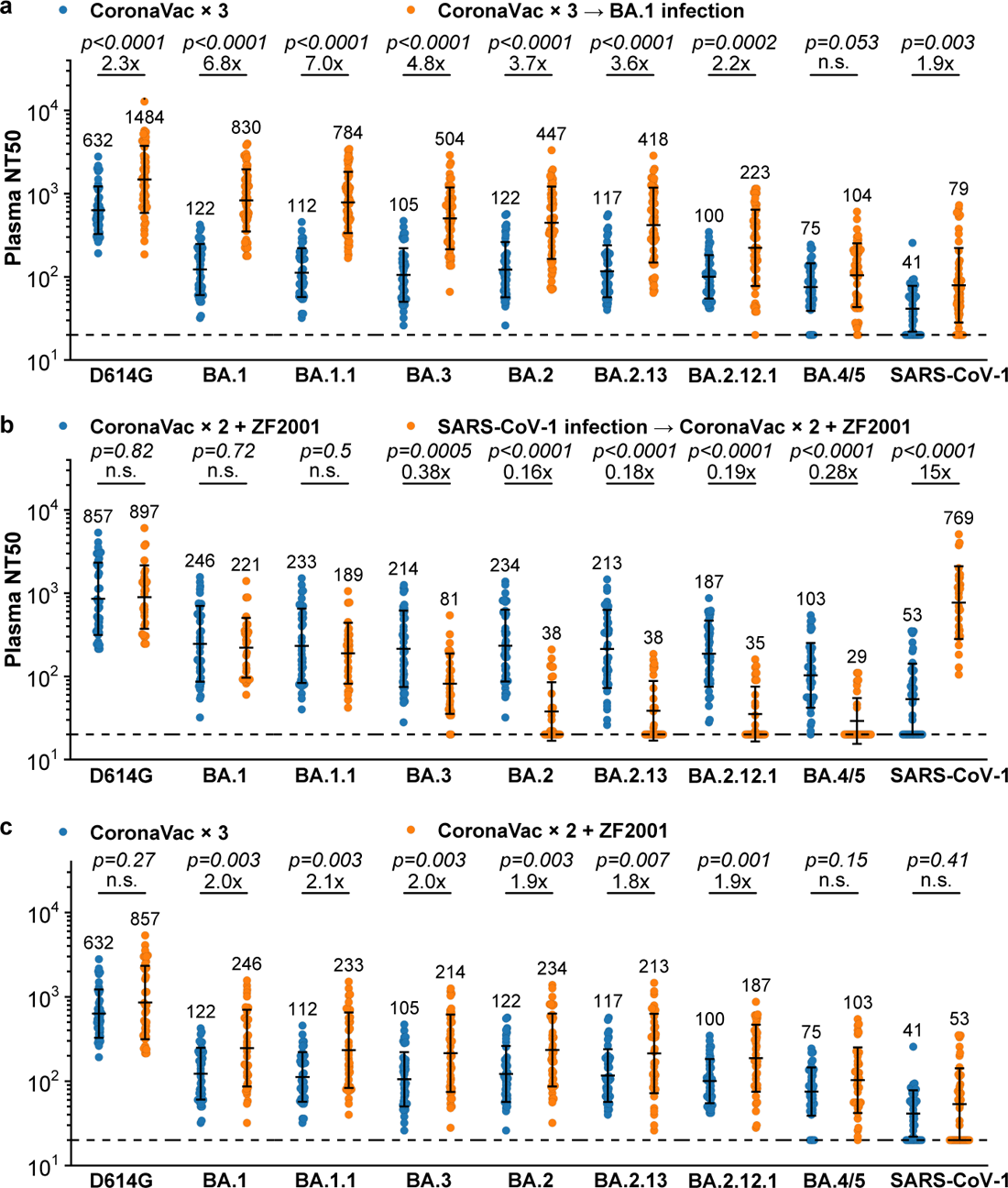
Different immunity backgrounds lead to distinct humoral immunity against Omicron subvariants NT50 against SARS-CoV-2, SARS-CoV-1 D614G and Omicron subvariants spike-pseudotyped VSV by plasma samples from. **a**, individuals who received 3 doses CoronaVac with (n=50) or without (n=40) BA.1 breakthrough infection; **b**, individuals who received 2 doses CoronaVac and ZF2001 booster with (n=28) or without (n=38) previous SARS-CoV-1 infection; **c**, individuals who received 3 doses CoronaVac (n=40) or 2 doses CoronaVac with ZF2001 booster (n=38). P-values were calculated using two-tailed Wilcoxon rank-sum tests and labeled above the bars. n.s., not significant, p>0.05. All neutralization assays were conducted in biological duplicates. Geometric means are labeled. Error bars refer to geometric standard deviations.

**Extended Data Fig. 3.**
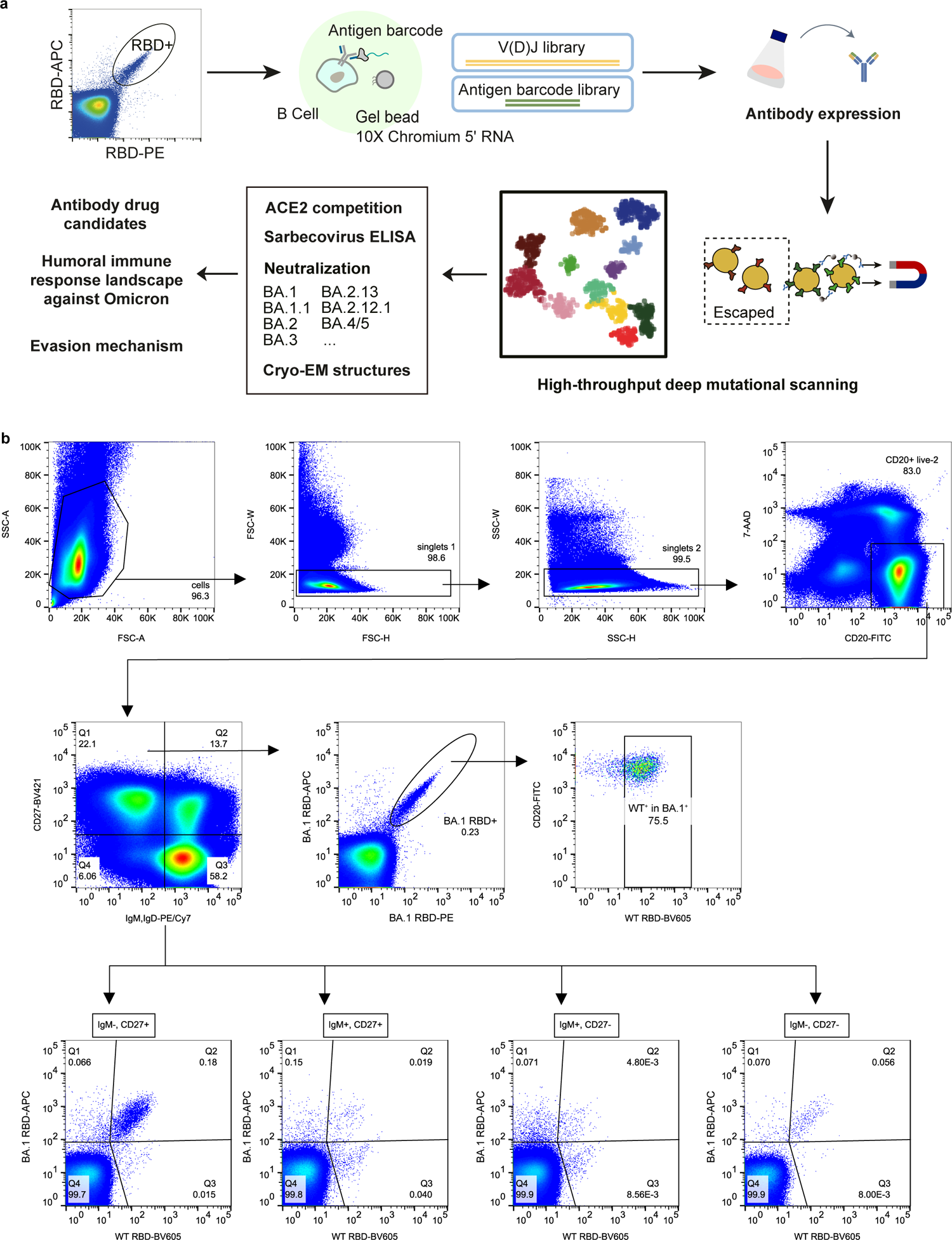
Workflow for the isolation and characterization of SARS-CoV-2 RBD antibodies. **a**, Overall schematic of antibody identification by single cell VDJ sequencing with feature barcodes and epitope analysis by high-throughput deep mutational scanning. **b**, FACS strategy to enrich BA.1/WT cross-reactive memory B cells or BA.1-specific memory B cells.

**Extended Data Fig. 4.**
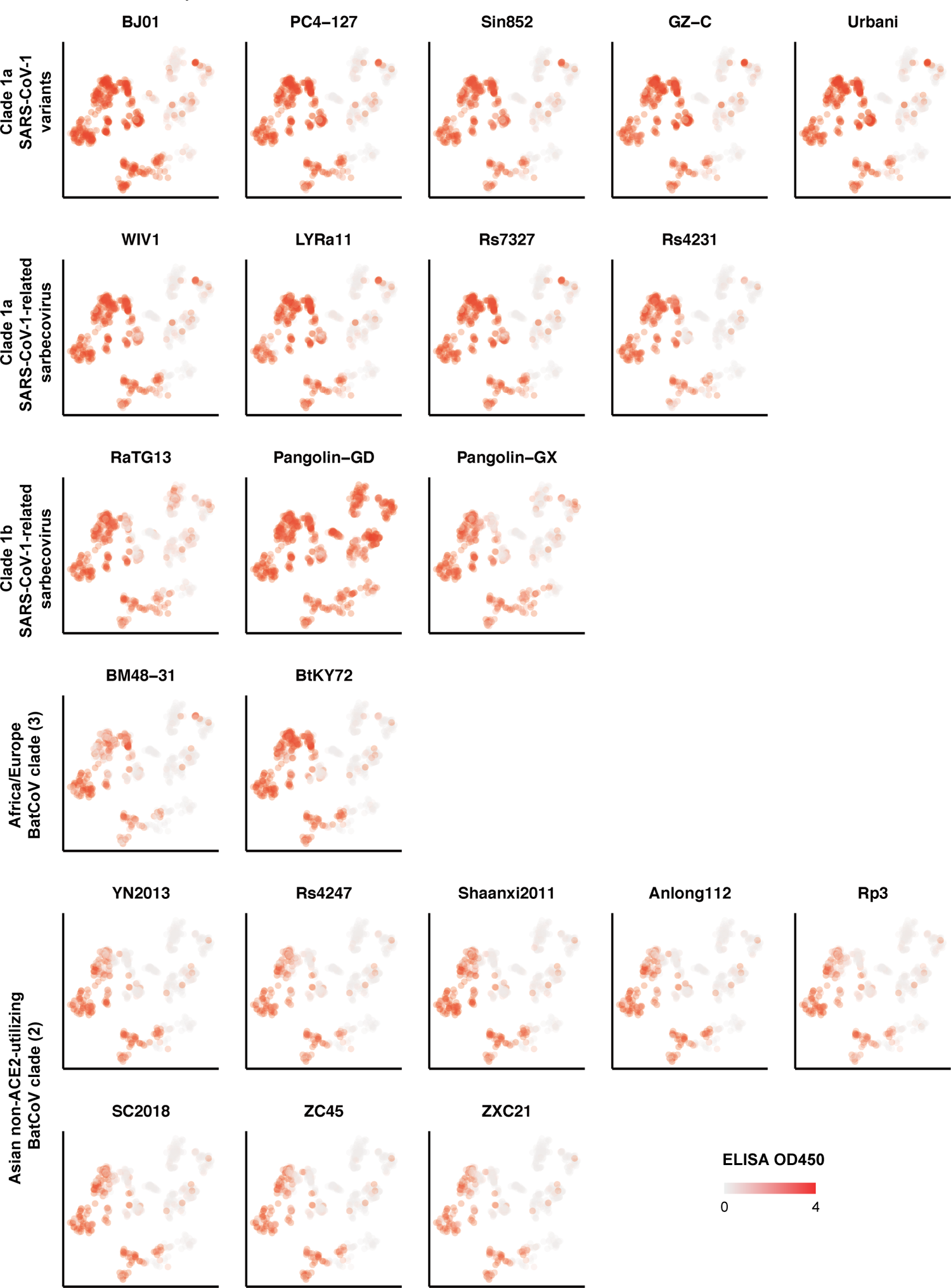
ELISA reactivity against 22 sarbecovirus RBD. Shades of red indicate ELISA OD450 for each antibody against various sarbecoviruses from different clades.

**Extended Data Fig. 5.**
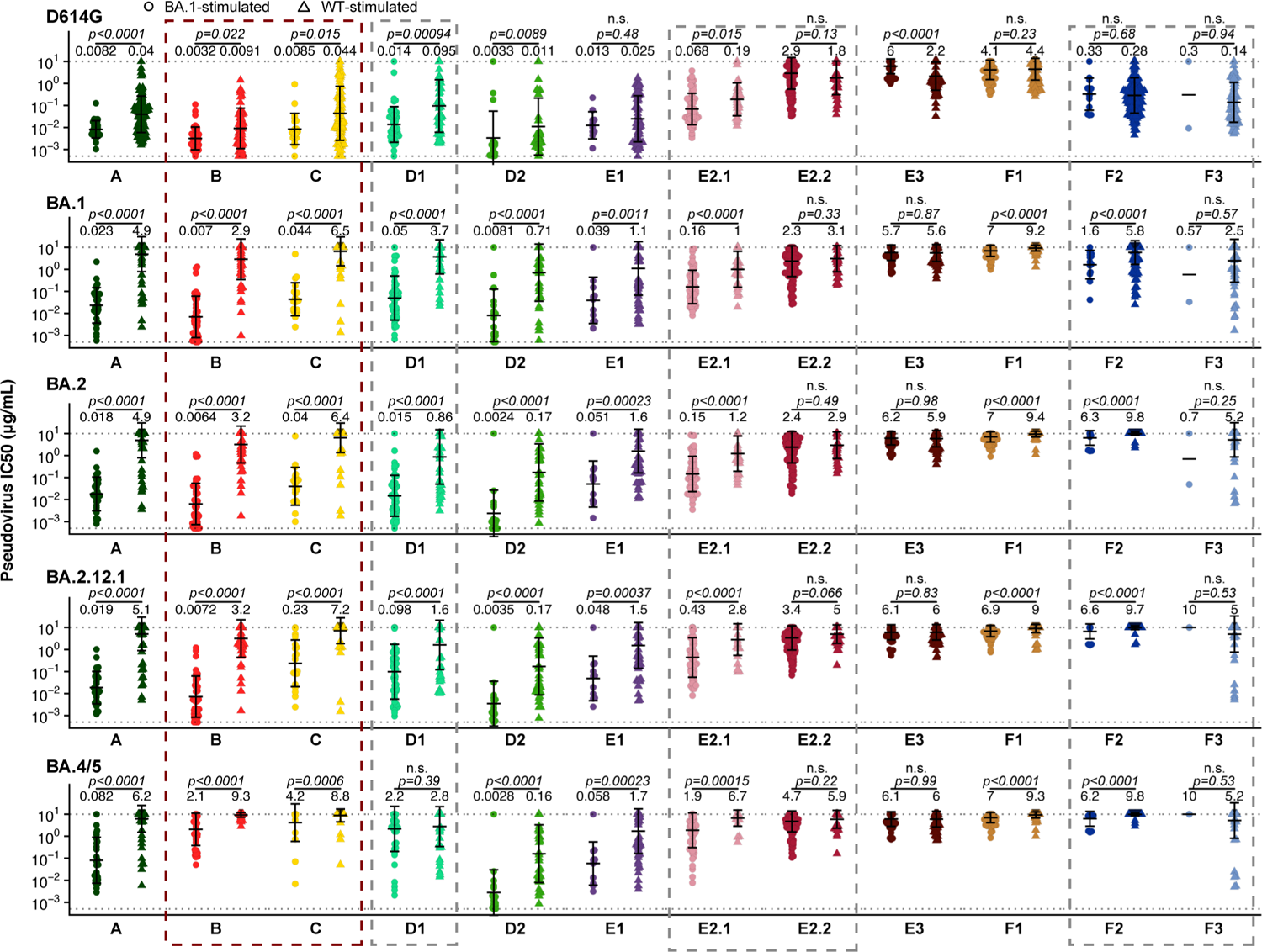
Neutralizing activities of antibodies elicited by SARS-CoV-2 BA.1 or wildtype. Neutralizing activity against SARS-CoV-2 D614G and Omicron subvariants pseudovirus by antibodies of each epitope group from BA.1 convalescents (BA.1-stimulated. A, n=30; B, n=41; C, n=20; D1, n=49; D2, n=17; E1, n=11; E2.1, n=64; E2.2, n=122; E3, n=57; F1, n=80; F2, n=13; F3, n=2), and from wildtype convalescents or vaccinees (WT-stimulated. A, n=98; B, n=55; C, n=88; D1, n=46; D2, n=36; E1, n=59; E2.1, n=26; E2.2, n=39; E3, n=68; F1, n=97; F2, n=158; F3, n=67). Geometric mean titers (GMT) are annotated above each group of points, and error bars indicate geometric standard deviation. P-values were calculated using two-tailed Wilcoxon rank-sum tests and labeled above the bars. n.s., not significant, p>0.05. NAbs in the boxed epitope groups showed substantial neutralization potency changes against BA.2.12.1 or BA.4/5 compared to BA.1. All neutralization assays were conducted in biological duplicates.

**Extended Data Fig. 6.**
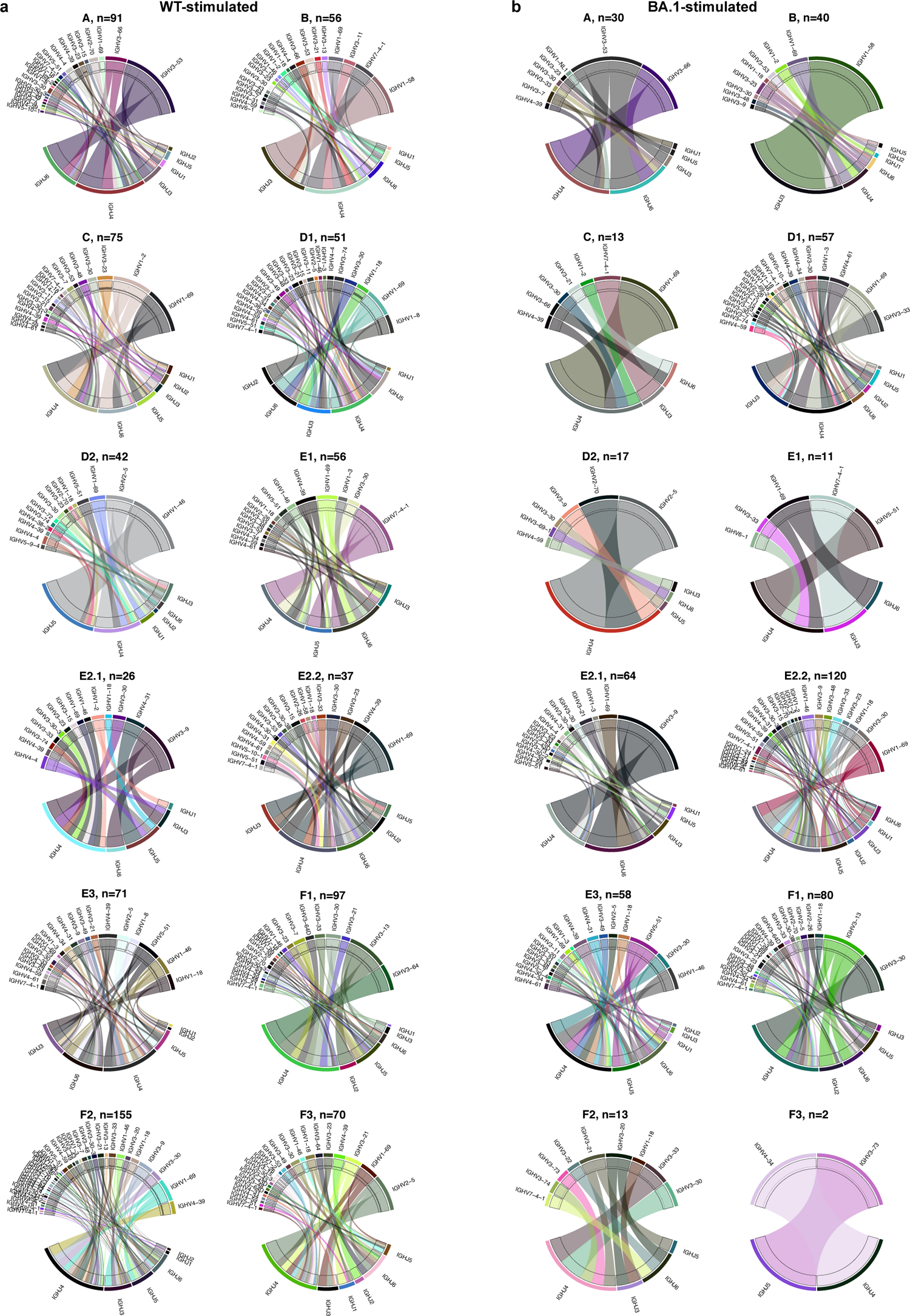
Heavy chain V-J genes of BA.1-stimulated and WT-stimulated antibodies in each epitope group. Heavy chain V-J genes combination of **a**, WT-stimulated antibodies. **b**, BA.1-stimulated antibodies or each epitope group. The number of NAbs is annotated above the chord plot.

**Extended Data Fig. 7.**
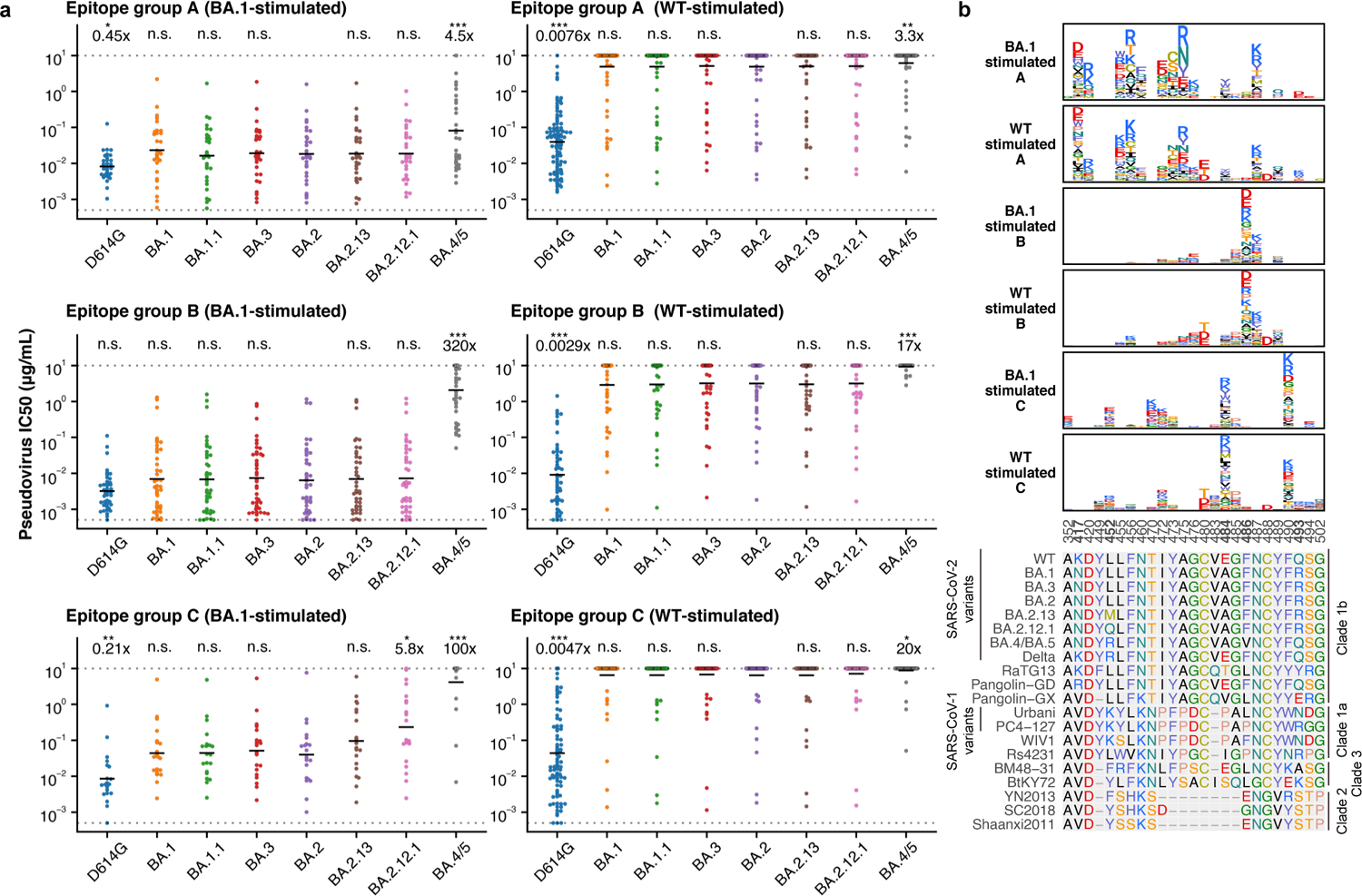
Comparison of BA.1-stimulated and WT-stimulated antibodies in group A, B and C. **a**, Neutralizing activity against SARS-CoV-2 D614G and Omicron subvariants by BA.1-stimulated (A, n=30; B, n=41; C, n=20) and WT-stimulated (A, n=98; B, n=55; C, n=88) antibodies in Group A, B and C. Geometric mean of IC50 fold changes compared to IC50 against BA.2 are annotated above the bars. P-values were calculated using a two-tailed Wilcoxon signed-rank test of paired samples, in comparison to IC50 against BA.2. *, p<0.05; **, p<0.01; ***, p<0.001; n.s., not significant, p>0.05. All neutralization assays were conducted in biological duplicates. **b**, Averaged escape maps at escape hotspots of BA.1-stimulated and WT-stimulated antibodies in group A, B and C, and corresponding MSA of various sarbecovirus RBDs. Height of each amino acid in the escape maps represents its mutation escape score. Mutated sites in Omicron variants are marked in bold.

**Extended Data Fig. 8.**
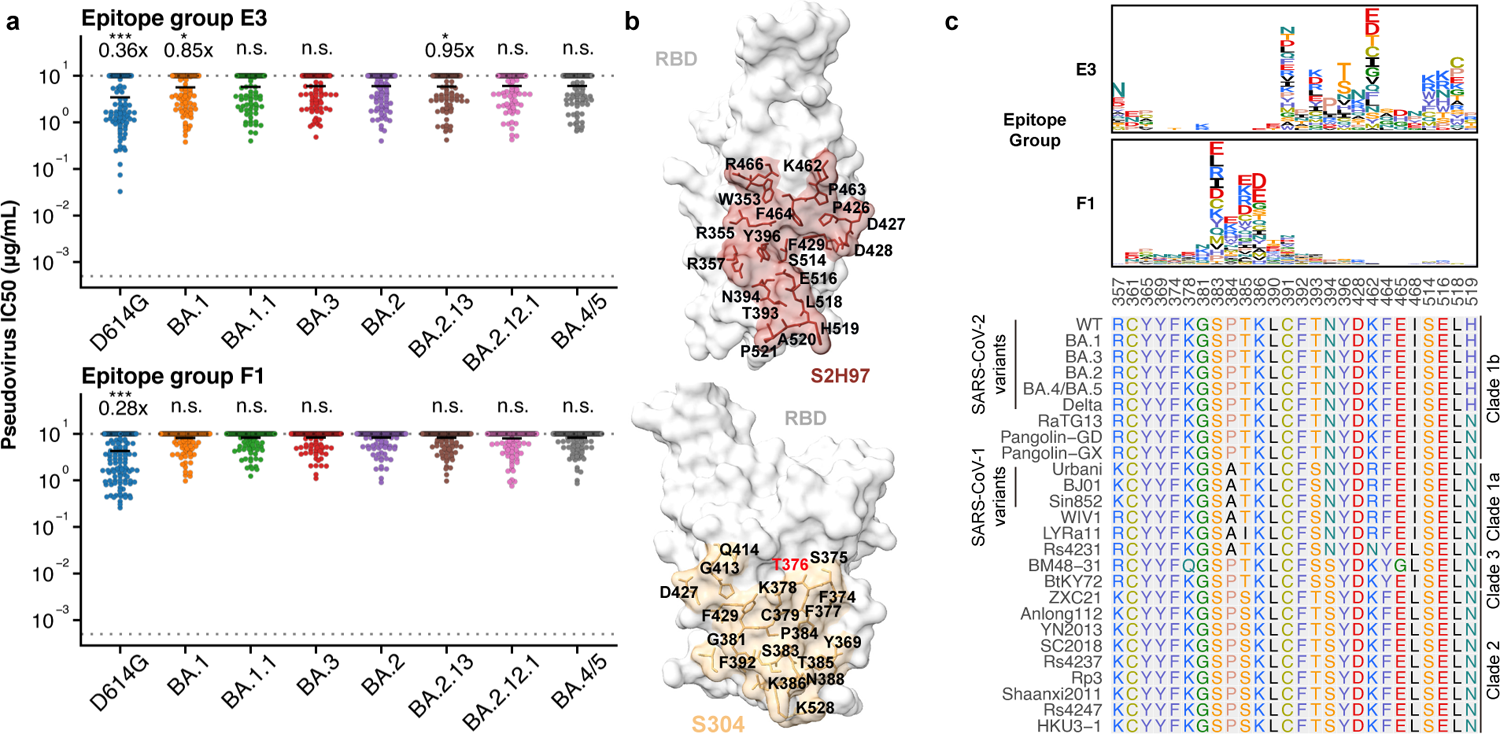
Antibodies of group E3 and F1 exhibit weak but broad-spectrum neutralization. **a**, Neutralizing activity against SARS-CoV-2 D614G and Omicron subvariants by antibodies in group E3 (n=125) and F1 (n=177). Geometric mean of IC50 fold changes compared to BA.2 are annotated above the bars. P-values were calculated using a two-tailed Wilcoxon signed-rank test of paired samples, in comparison to IC50 against BA.2. *, p<0.05; **, p<0.01; ***, p<0.001; n.s., not significant, p>0.05. All neutralization assays were conducted in biological duplicates. **b**, Epitope of representative antibodies in group E3 (S2H97, PDB: 7M7W) and F1 (S304, PDB: 7JW0). Residues highlighted in red indicate mutated sites in Omicron variants. **c**, Averaged escape maps at escape hotspots of antibodies in group E3 and F1, and corresponding MSA of various sarbecovirus RBDs. Height of each amino acid in the escape maps represents its mutation escape score. Mutated sites in Omicron variants are marked in bold.

**Extended Data Fig. 9.**
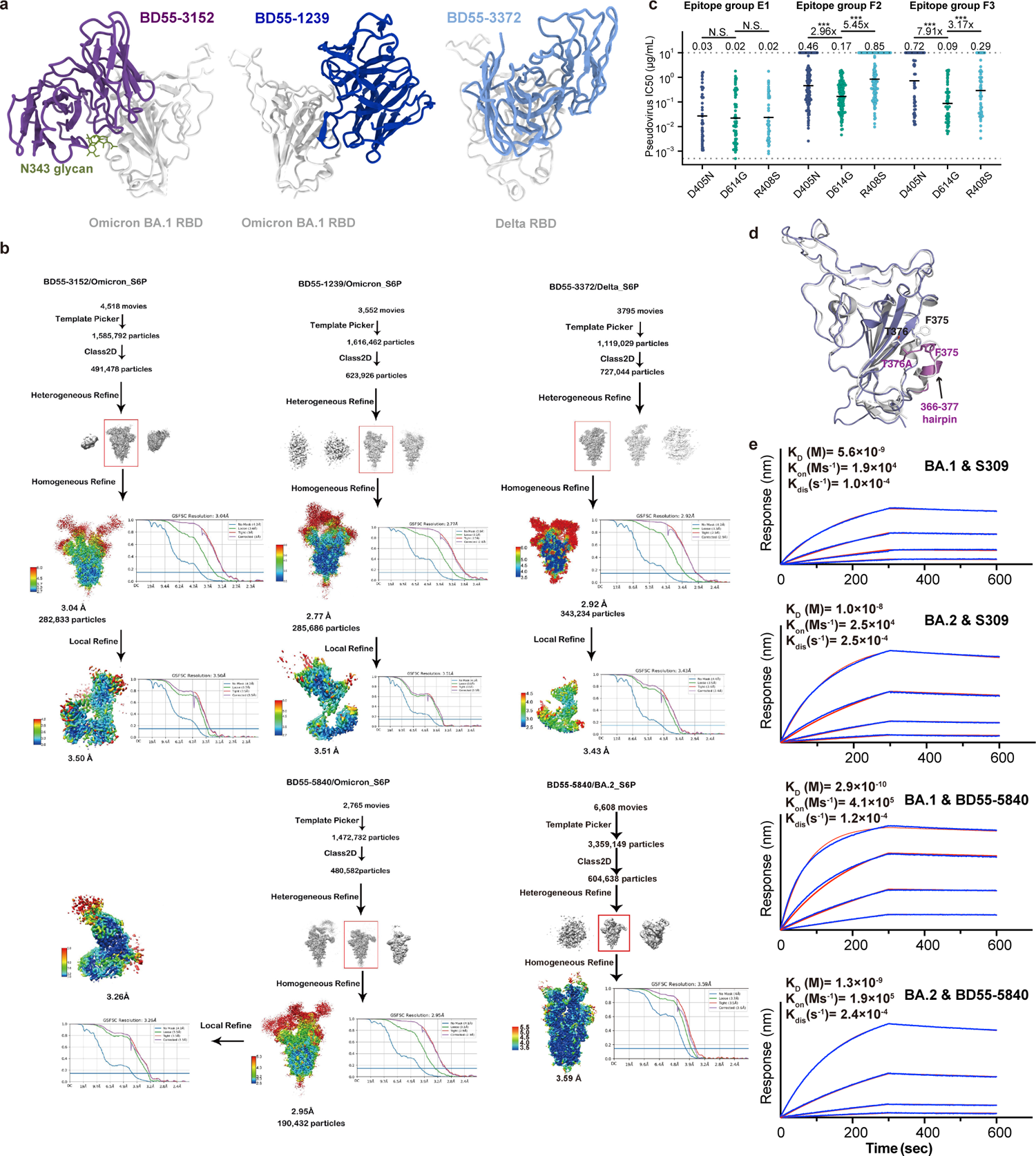
RBD-binding structures and affinity of broad Sarbecovirus antibodies. **a**, Cartoon models of Cryo-EM structures of BD55-3152 in complex of BA.1 RBD, BD55-1239 in complex of BA.1 RBD, and BD55-3372 in complex of Delta RBD. **b**, Workflow to generate refined structural model of BD55-3152 and BD55-1239 in complex of BA.1 RBD, BD55-3372 in complex of Delta RBD, and BD55-5840 in complex of BA.2 RBD. **c**, Neutralizing activity of representative NAbs in group E1 (n=68), F2 (n=139) and F3 (n=61) against SARS-CoV-2 D614G, in addition to D614G+D405N and D614G+R408S. Geometric mean of IC50 fold changes compared to IC50 against D614G are annotated above the bars. P-values were calculated using a two-tailed Wilcoxon signed-rank test of paired samples. *, p<0.05; **, p<0.01; ***, p<0.001; n.s., not significant, p>0.05. All neutralization assays were conducted in biological duplicates. **d**, Conformational comparison between BA.1 and BA.2 RBD regarding the 366-377 hairpin. **e**, Biolayer interferometry analysis of Group E1 antibodies S309 and BD55-5840 binding to Omicron BA.1 and BA.2 Spike trimer. Biolayer interferometry analyses were conducted in biological duplicates.

**Extended Data Fig. 10.**
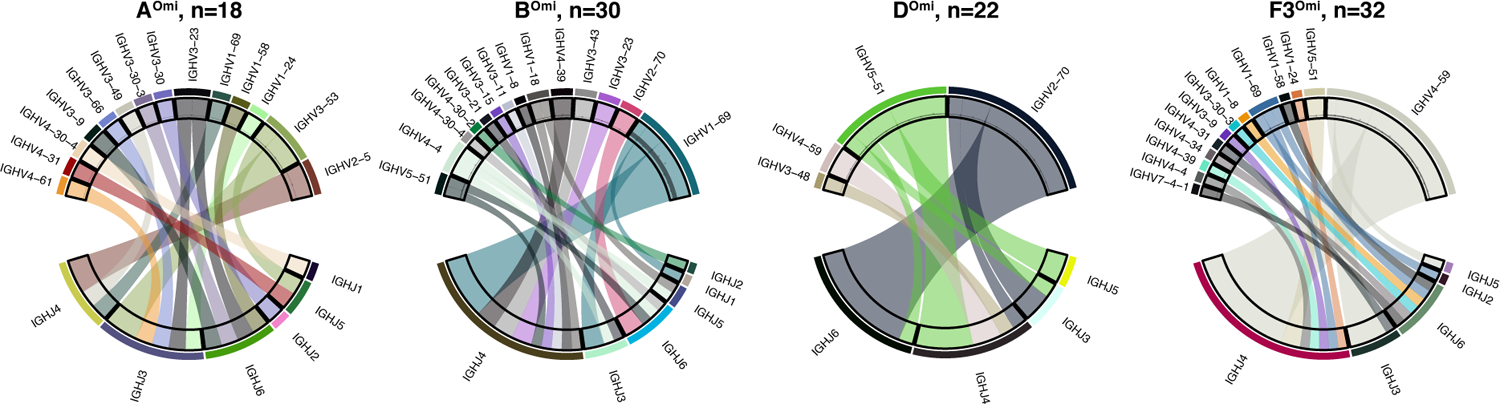
HV-HJ gene combination of BA.1-specific antibodies. Heavy chain V-J gene combination of BA.1-specific neutralizing antibodies in BA.1-specific epitope groups A^Omi^, B^Omi^, D^Omi^ and F3^Omi^.

