## Supplementary Guide for "BA.2.12.1, BA.4 and BA.5 escape antibodies elicited by Omicron infection"

**Supplementary Information Guide**

**Supplementary Table 1**

Summarized information of SARS-CoV-2 vaccinated individuals, BA.1 convalescents and SARS convalescents involved in the study.

**Supplementary Table 2**

Summarized information and experiment results of 1640 SARS-CoV-2 RBD antibodies involved in this study, including their sources, epitope groups, pseudovirus neutralizing IC50 and ELISA OD450 for sarbecovirus, ACE2 competition levels, and heavy/light chain sequences.

**Supplementary Table 3**

Accession numbers of sequences of sarbecovirus RBD used in the study.

**Supplementary Table 4**

Reports of detailed parameters of the reconstructed Cryo-EM structures.
